# Redundant Trojan horse and endothelial-circulatory mechanisms for host-mediated spread of *Candida albicans* yeast

**DOI:** 10.1101/2020.02.21.959387

**Authors:** Allison K. Scherer, Bailey A. Blair, Jieun Park, Brittany G. Seman, Joshua B. Kelley, Robert T. Wheeler

## Abstract

The host innate immune system has developed elegant processes for the detection and clearance of invasive fungal pathogens. These strategies may also aid in the spread of pathogens *in vivo*, although technical limitations have previously hindered our ability to view the host innate immune and endothelial cells to probe their roles in spreading disease. Here, we have leveraged zebrafish larvae as a model to view the interactions of these host processes with the fungal pathogen *Candida albicans in vivo*. We examined three potential host-mediated mechanisms of fungal spread: movement inside phagocytes in a “Trojan Horse” mechanism, inflammation-assisted spread, and endothelial barrier passage. Utilizing both chemical and genetic tools, we systematically tested the loss of neutrophils and macrophages and the loss of blood flow on yeast cell spread. Both neutrophils and macrophages respond to yeast-locked and wild type *C. albicans* in our model and time-lapse imaging revealed that macrophages can support yeast spread in a “Trojan Horse” mechanism. Surprisingly, loss of immune cells or inflammation does not alter dissemination dynamics. On the other hand, when blood flow is blocked, yeast can cross into blood vessels but they are limited in how far they travel. Blockade of both phagocytes and circulation reduces rates of dissemination and significantly limits fungal spread distance from the infection site. Together, this data suggests a redundant two-step process whereby (1) yeast cross the endothelium inside phagocytes or via direct uptake, and then (2) they utilize blood flow or phagocytes to travel to distant sites.

**Author summary:** Although *Candida albicans* is the most common cause of fungal bloodstream infection, we know little about how it spreads from one tissue to another. Host processes often inadvertently assist pathogens that can hijack immune cells or induce endothelial endocytosis. Here we have used transparent zebrafish to visualize how specific host cells and processes contribute to yeast spread *in vivo.* We find that *Candida* is a sophisticated pathogen that uses redundant strategies to disseminate in the host: it can either use host immune cells or cross into the circulatory system and use blood flow. These data suggest that both of these mechanisms must be targeted to limit *Candida* dissemination during infection.

## Introduction

*Candida albicans* is a small non-motile fungus that can cause disseminated candidiasis in immunocompromised populations. This dimorphic pathogen is a normal commensal organism of the mouth, gastrointestinal track, and vagina that causes life-threatening invasive candidiasis in immunocompromised patients. *C. albicans* is able to spread from mucosal sites to internal organs in immunocompromised mouse disease models and this is believed to be a primary infection route for this fungus in humans as well [1–4]. Recent work using a tissue-to-bloodstream *C. albicans* dissemination model in zebrafish has established that the yeast form is specialized for promoting infection spread [5]. Although we still do not understand how yeast orchestrate this movement, it has been proposed that *C. albicans* spreads via phagocyte-dependent or –independent mechanisms [6, 7].

Bacterial and fungal pathogens utilize the host’s immune cells in a “Trojan Horse” mechanism to spread to outlying tissues, stimulating engulfment, surviving within the phagocyte for sufficient time to allow migration away from the infection, then provoking release from the phagocyte. Both *in vitro* and zebrafish disease models have been used to demonstrate how neutrophils [8] and macrophages [9–12] can be vehicles for dissemination for mycobacterium, *Cryptococcus* and *Streptococcus*. Although *C. albicans* was previously suspected of being an extracellular pathogen based on *in vitro* challenges, intravital imaging in the zebrafish model has suggested that it can establish an impasse with macrophages that is a prerequisite for migration from the infection site [13].

Independently of phagocyte-driven spread, *C. albicans* may be moved directly into the bloodstream from tissues via endocytosis by endothelial cells or paracellular invasion. *C. albicans* can cause endothelial cell damage and stimulate endocytosis *in vitro*, suggesting one strategy for invasion through this barrier [3, 7, 14, 15]. Once fungi get into the bloodstream, they can be carried by the circulatory system throughout the host, a process that results in rapid and universal dispersion [16]. Movement of *C. albicans* yeast through the endothelial barrier has not yet been studied *in vivo*, largely due to technical limitations in murine models.

To systematically test the importance of these host-mediated mechanisms of yeast spread for *C. albicans* infection, we used a recently described zebrafish yolk infection model [5, 17]. This anatomical structure represents a simplified system where translocation through the endothelium and spread throughout the bloodstream can be monitored through longitudinal imaging of both fluorescent fungi and phagocytes. Using high-resolution intravital imaging, we established that both macrophages and neutrophils engulf yeast, while *C. albicans* survives within macrophages and can be released far from the site of infection through non-lytic exocytosis. Systematic elimination of both macrophage and neutrophil function, however, resulted in no loss in dissemination frequency or levels. Similarly, elimination of blood flow did not in and of itself result in a reduction in dissemination. When both phagocytes and blood flow were disabled, however, there was a significant decrease in levels of dissemination. Thus, phagocyte-dependent and –independent modes are redundant and the versatility of *C. albicans* in using different strategies ensures robust spread of infection.

## Results

### *C. albicans* yeast dissemination from localized infection is preceded by innate immune cell recruitment

Pathogen spread within a host is crucial for pathogenesis, but understanding the role of the host in this process has been particularly difficult to quantify in mammalian infection models. Fungal pathogens can disseminate as yeast or hyphae, and recent work using a larval model of zebrafish infection suggests that yeast are specialized for spread from tissue to bloodstream. We sought to use this uniquely tractable model to probe the host’s role in yeast dissemination, postulating that yeast may spread through phagocyte-dependent or –independent strategies.

We first sought to determine if yeast dissemination is passive or driven by specific fungal-specific determinants. We reasoned that if dissemination were passive it might be triggered by a given level of fungal burden, enhanced by proximity to the vasculature, or even occur with dead yeast. Previous work has shown that different starting inoculums of yeast don’t influence dissemination rate [5], which suggested that dissemination frequency is relatively insensitive to inoculum size. We also found no difference in fungal burden among groups of recruited and disseminated larvae (Fig. S1A-C), suggesting that neither dissemination nor phagocyte responses are driven by the presence of a threshold of NRG1^OEX^. Comparing the location of yeast in the yolk with distance to fluorescent vascular endothelium, we found that proximity of the infection site to blood vessels is not associated with spread of infection (Fig. S2A-D). Finally, we found that dissemination is significantly reduced with killed yeast, suggesting that yeast provide essential cues to mediate their own spread (Fig. S3A-B). Together, these data suggest that dissemination and immune recruitment are driven by active rather than passive processes.

To understand if host innate immune responses are associated with *C. albicans* spread, we infected transgenic zebrafish larvae in the yolk with yeast-locked fungi and intravitally imaged both host immune cells and fungi. Larvae were infected with yeast-locked NRG1^OEX^ (NRG1^OEX^-iRFP; Fig 1A) and scored for recruitment of phagocytes to the infection and yeast cell spread from the infection site. Fish were almost exclusively found to be in only three of the expected four classes, with about 50% in the “No Recruitment/No Dissemination” class, almost none in the “No Recruitment/Dissemination” class, and 25% each in the other two classes (Fig. 1B). To better understand if immune involvement was associated with yeast dissemination or infection resolution, larvae were longitudinally imaged over a 16-hour time course. Larvae that had recruited macrophages at 24 hpi stayed the same or progressed to a recruited state with NRG1^OEX^-iRFP dissemination at 40 hpi. Larvae that began with recruited macrophages and disseminated NRG1^OEX^- iRFP did not revert to a non-recruited or non-disseminated state. In limited cases, dissemination occurred without the response of phagocytes to the yolk sac, and only in one case did it lead to later phagocyte recruitment to the yolk sac. These experiments demonstrated that recruitment of phagocytes tends to precede dissemination (Fig. 1B-C), suggesting a role for phagocytes in the dissemination of yeast, either directly through the Trojan Horse mechanism, or indirectly through inflammation-mediated weakening of host vasculature [12, 18, 19].

**Figure 1.**
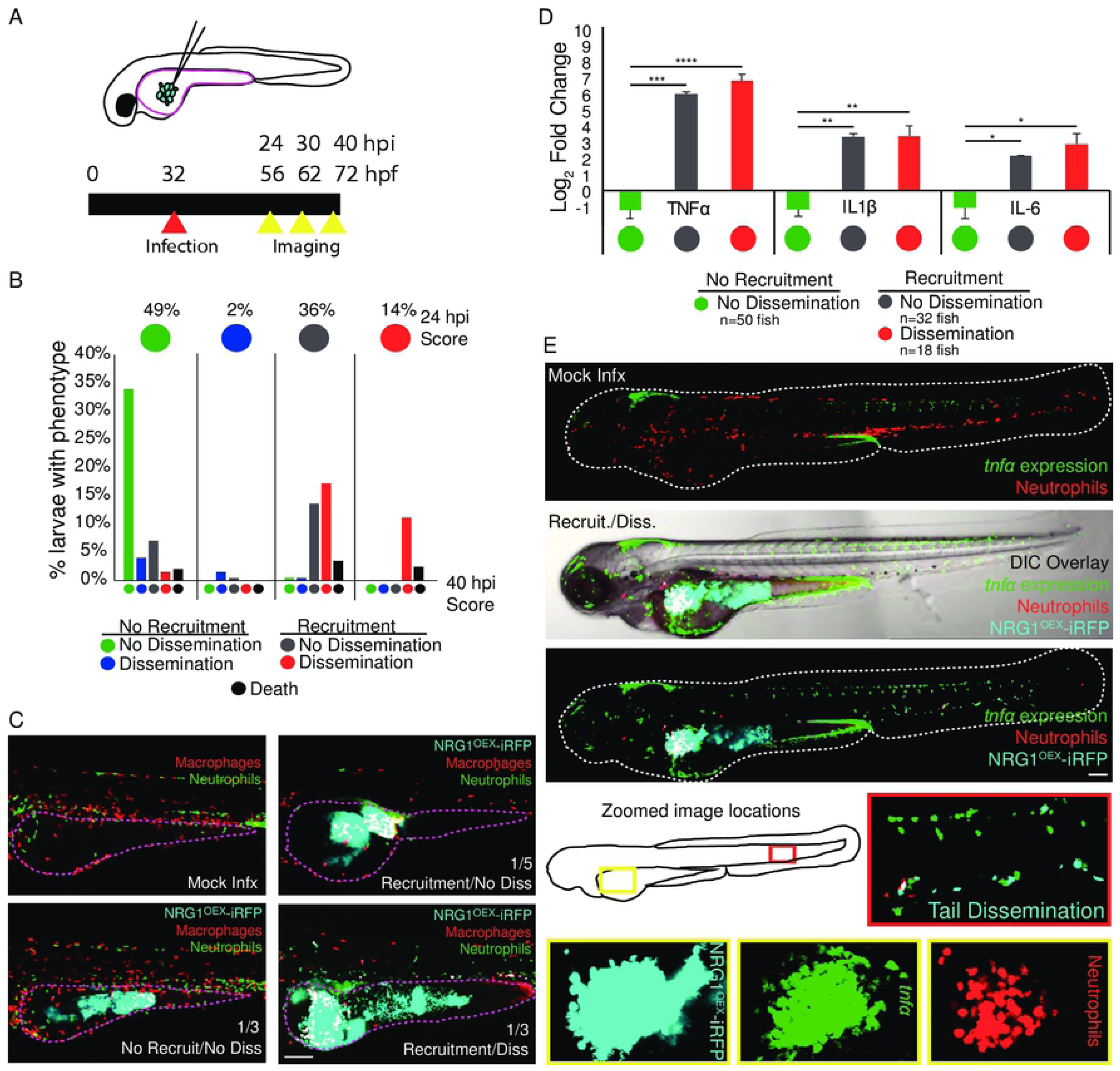
Macrophage and neutrophil recruitment to the site of infection is positively associated with dissemination and proinflammatory cytokine upregulation. Larvae were infected with far-red fluorescent yeast-locked *C. albicans NRG1*^OEX^-iRFP in the yolk sac at ∼32 hours post fertilization (hpf). (A) Schematic of the yolk sac injection and timeline of the infection and screening for both recruitment of phagocytes and dissemination of yeast. (B) Progression of infection in which larvae with early recruitment of macrophages go on to develop disseminated infection and larvae rarely revert back to a non-disseminated state. Top circles represent the percentages of fish at 24 hpi, colored by recruitment/dissemination phenotype. Bottom bars represent subsequent phenotypes at 40 hpi, grouped based on their 24 hpi phenotype. Thus, 49% of fish at 24 hpi had the NR/ND phenotype (green circle at top left). Considering both 24 hpi and 40 hpi scores, 33% of fish were in the category: NR/ND at 24 hpi and NR/ND at 40 hpi (green bar on far left). Scores were pooled from larvae with fluorescent macrophages from a total of 12 experiments: 5 experiments in *Tg(mpeg:GAL4)/(UAS:Kaede)*, 4 experiments in *Tg(mpeg:GAL4)/(UAS:nfsb-mCherry)*, and 3 experiments in *Tg(fli1:EGFP)* x *Tg(mpeg:GAL4)/(UAS:nfsb-mCherry)*. A total of 197 fish were followed from 24 to 40 hpi. (C) Examples of each infection category in *Tg(mpeg:GAL4)/(UAS:nfsb-mCherry)/Tg(mpx:EGFP)* larvae. The yolk sac is outlined in a dotted magenta line and the approximate proportion of larvae with this score at 40 hpi is indicated in the lower right of the image (the remainder of fish died between 24 and 40 hpi). Scale bar = 150 µm. (D) *Tg(mpeg:GAL4)/(UAS:nfsb-mCherry)* larvae were scored for macrophage recruitment to the site of infection and yeast dissemination at 24 hpi, then groups of 6-10 were homogenized for qPCR. Gene expression for TNFα, IL1β, and IL-6 were measured and mock-infected larvae were used for reference. Data come from 3 independent experiments with 97 PBS larvae, 50 No recruitment/No dissemination, 32 Recruited/No dissemination, and 18 Recruited/Disseminated total larvae used for analysis. Means are shown with standard error of measurement. One-way ANOVA with Dunnett’s multiple-comparisons post-test demonstrates significantly higher gene expression in larvae with recruited macrophages at the site of infection, regardless of dissemination scores (* p≤ 0.05, ** p ≤ 0.01, *** p ≤ 0.001, **** p < 0.0001). (E) Representative *Tg(lysC:Ds-Red)/Tg(tnfa:GFP)* larvae with red fluorescent neutrophils and green fluorescence with *tnfα* expression in either a mock infected larvae (top) or a yeast-locked *C. albicans* infected fish. Neutrophils are recruited to the yolk sac and *tnfα* is expressed at the site of infection and in the tail with disseminated yeast cells. Scale bar = 100 µm.

### Phagocyte recruitment correlates with pro-inflammatory gene upregulation and local expression of TNFα

Phagocytes can indirectly facilitate pathogen dissemination by proteolysis of epithelial cell junctions and by producing pro-inflammatory cytokines that increase local vascular permeability [20–23]. We therefore examined if dissemination events were associated with increases in inflammatory cytokine production. We scored for infection spread and recruitment of macrophages and measured cytokine mRNA levels in pooled larvae from each infection category. Larvae that had recruitment of macrophages to the site of infection had significant increases for TNFα, interleukin-1-beta (IL1β), and interleukin-6 (IL-6), and this increase in expression in recruited larvae was independent of dissemination (Fig. 1D). We found similar results for infected larvae with both fluorescent neutrophils and macrophages, where increases in TNFα, IL1β, and IL-6 were also independent of dissemination (data not shown). These data suggest that inflammatory gene expression is associated with phagocyte recruitment, and that cytokine expression at the infection site might precede dissemination.

As TNFα was found to be upregulated in all larvae with phagocyte recruitment scores, we were interested in determining where TNFα was being expressed in relation to the location of yeast. A transgenic reporter zebrafish line of *tnfa* transcriptional activity was used to visualize the expression of TNFα in *C. albicans* infected fish [22]. We found *tnfa:EGFP* expression localized primarily at the site of infection and occasionally in phagocytes interacting with disseminated fungi far from the infection site (Fig. 1E). Consistent with the qPCR results, *tnfa:EGFP* expression at the site of infection tended to correlate with phagocyte recruitment but not with dissemination (Fig. S4). Taken together, these data suggest that phagocytes recruited to the site of infection produce locally high levels of proinflammatory cytokines that may alter the infection environment, but elevated cytokine levels are not sufficient to drive dissemination events.

### Innate immune cells transport yeast into and throughout the bloodstream

Phagocytes unwittingly spread viruses, intracellular bacteria and intracellular fungi [12, 24–28]. Until recent intravital imaging experiments, it was not appreciated that *C. albicans* can survive inside phagocytes and create an impasse that could potentially abet its spread into new host tissues through a Trojan Horse mechanism [13]. This “Trojan Horse” mechanism has three distinct stages: first, phagocytes respond to the infection and engulf the pathogen, then phagocytes move out of the infection site, and finally the pathogen exits the phagocyte in distant tissues. To look at overall phagocyte-fungi interaction dynamics in yolk infection, we quantified the number of neutrophils and macrophages at the initial infection site over time. We found that, for fish with fungal dissemination and phagocyte recruitment at 24 hpi, there was an overall decrease in the number of phagocytes at the site of infection over the next 16 hrs (Fig. S5A-B; statistically significant difference for macrophages and consistent trend for neutrophils). There was no overall trend for fish that did not have recruitment and dissemination at 24 hpi. This decrease in phagocyte amount at the later time point was either because immune cells were leaving the site or dying as a result of infection [29], but time-course images couldn’t distinguish between these possibilities. Therefore, several time-lapse experiments were performed to examine the fate of macrophages with internalized yeast. These show that macrophages do phagocytose and carry yeast away from the site of infection and into the yolk sac circulation valley (Fig. 2A, Movies S1 and S2).

**Figure 2.**
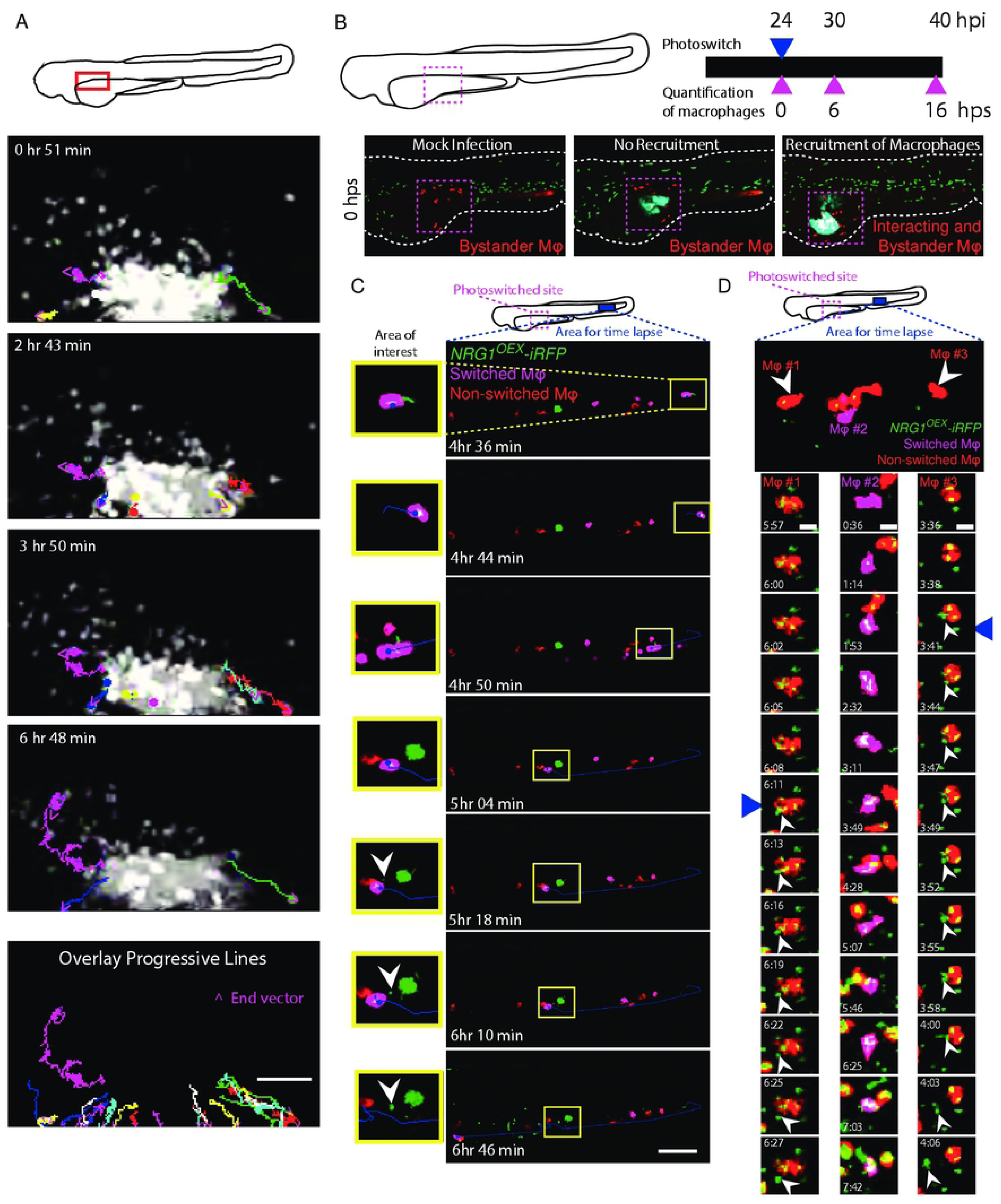
Phagocytes actively participate in the transport of yeast within the bloodstream. *Tg(mpeg1:GAL4/UAS:nfsb-mCherry)/Tg(mpx:EGFP)* or *Tg(mpeg1:GAL4)x(UAS:Kaede)* larvae were infected with yeast-locked *C. albicans* and scored for recruitment of macrophages and dissemination of yeast at 24 hpi. (A) Time-lapse taken over the course of 7 hours starting at ∼32 hpi. The schematic above illustrates the area imaged. Neutrophils and macrophages were observed moving within and away from the infection site carrying yeast. Dots and lines indicate tracks taken by phagocytes during the time lapse, with magenta arrowheads indicating the direction of movement. Phagocytes were observed taking yeast from the site of infection in these cases. The bottom panel illustrates all tracked paths from the time lapse in a single overlay image. Scale bar = 50 µm. Frames taken from Movie S1. (B) A schematic of our macrophage photo-conversion experiments where *Tg(mpeg1GAL4) x Tg(UAS:Kaede)* macrophages near the infection site were photoswitched at 24 hpi, and confocal images of each photoswitched larva were taken at 24, 30, and 40 hpi to track macrophages. Macrophages near the site of infection but not interacting with yeast were identified as “bystander” macrophages. (C) Time-lapse panels of *Tg(mpeg1GAL4) x Tg(UAS:Kaede)* larva with a photoconverted macrophage moving out of the blood stream into tail tissue. The top schematic shows the regions used for photoswitching (magenta dashed box) at 24 hpi and the region selected for time-lapse (magenta solid box) from ∼46 hpi to ∼53 hpi. The yellow highlighted area demonstrates a photo-converted macrophage stopping in the bloodstream, rolling down the tail, and then releasing yeast. White arrowhead in the image panel highlights an apparent non-lytic expulsion (NLE) event. Time stamps indicate HR:MIN. Frames taken from Movie S3. Scalebar = 100 µm. (D) (Top) Area of tail imaged by time-lapse microscopy in an independent fish from that shown in (C). White arrowheads indicate macrophages followed in detail below. (Below) Middle column of frames highlights yeast growth within a photo-converted macrophage and left and right columns of frames show apparent NLE release of yeast cells from macrophages. Time stamps indicate HR:MIN. White arrowheads within images indicate apparent NLE events. Frames are taken from Movie S4. Scalebar = 10 µm.

Taken together, we see that phagocytes respond to the infection, phagocytose yeast and carry them from the infection site, but this left open the question of if and how the pathogen escapes from the phagocyte. To follow macrophages from the infection site, we used a transgenic fish with photo-switchable Kaede-expressing macrophages. Previous work with this photoconvertible transgenic line showed that macrophages that phagocytose fungi tend to die at the infection site rather than move away [29], but we found that a minority of macrophages with phagocytosed yeast remained viable and migrated from the infection site (Fig. 2B). We followed photo-switched macrophages, imaged *C. albicans*-macrophage dynamics at 40 hpi in distant infection foci, and found instances of *C. albicans* being released from macrophages in each of two larvae within one experiment. In the first larvae, two photoconverted macrophages exocytosed three yeast (Fig. 2C and Movie S3) and in the second larvae one macrophage released a yeast (Fig. 2D and Movie S4). These time-lapses include clear instances of yeast escaping from macrophages far from the infection site, providing evidence that the third step of the process of phagocyte-aided dissemination does occur *in vivo*.

### Neutrophil inactivation coupled with macrophage ablation does not alter infection outcomes

While these time-lapse experiments documented the occurrence of Trojan Horse-mediated spread, they are anecdotal and the high-content nature of the experiments prevented quantification of these events in large numbers of infected fish of all cohorts to determine the relative contribution of infected macrophages to overall dissemination. To define the importance of phagocyte-mediated dissemination, we took advantage of the ability to ablate macrophages and disable neutrophils in the zebrafish larva.

We first investigated how much the ablation of macrophages alters yeast spread, eliminating macrophages either by injection of liposomal clodronate [9, 25, 30, 31] or by addition of metronidazole pro-toxin to larvae with nitroreductase-expressing macrophages [32–34]. We confirmed that both methods are effective at eliminating macrophages (Fig. S6). Larvae treated with clodronate liposomes or metronidazole had an almost complete loss of macrophages, and no change in recruited neutrophils (Fig. S6A-D). To our surprise, ablation of macrophages by either method did not result in any alteration in dissemination frequencies at 40 hpi (Fig. S7A). Furthermore, ablated larvae also had a similar progression of events where phagocyte recruitment (neutrophils) preceded fungal dissemination (Fig. S7B). Interestingly, there was a trend among disseminated fish towards scores of “High Dissemination” in macrophage ablated fish, when the number of disseminated cells was counted (1-10 = Low, 11-50 = Medium, >50 = High; p = 0.076 for clodronate liposomes and p = 0.103 for metronidazole; Fig. S7A). We complemented these chemical ablation experiments with additional experiments using *pu.1* morpholinos to block macrophage development. This method also yielded similar dissemination frequencies of yeast with and without macrophage development (Fig. S8). Taken together, these experiments, using three different ways of eliminating macrophages, clearly show that macrophages are unexpectedly not required for efficient dissemination.

Because we observed neutrophils at the infection site and also interacting with disseminated yeast in the tail (Movies S3-S5), we sought to determine if neutrophils could substitute for macrophages in transporting yeast. We used Rac2-D57N larvae, with defective neutrophils that are unable to extravasate from the bloodstream [35], and eliminated macrophages with clodronate. To our surprise, there was again no difference in dissemination rates in the context of impaired neutrophils, ablated macrophages, or both (Fig. 3A). The fact that elimination of both major phagocyte types does not alter dissemination infection dynamics suggests that Trojan Horse infection spread is a redundant mechanism.

**Figure 3.**
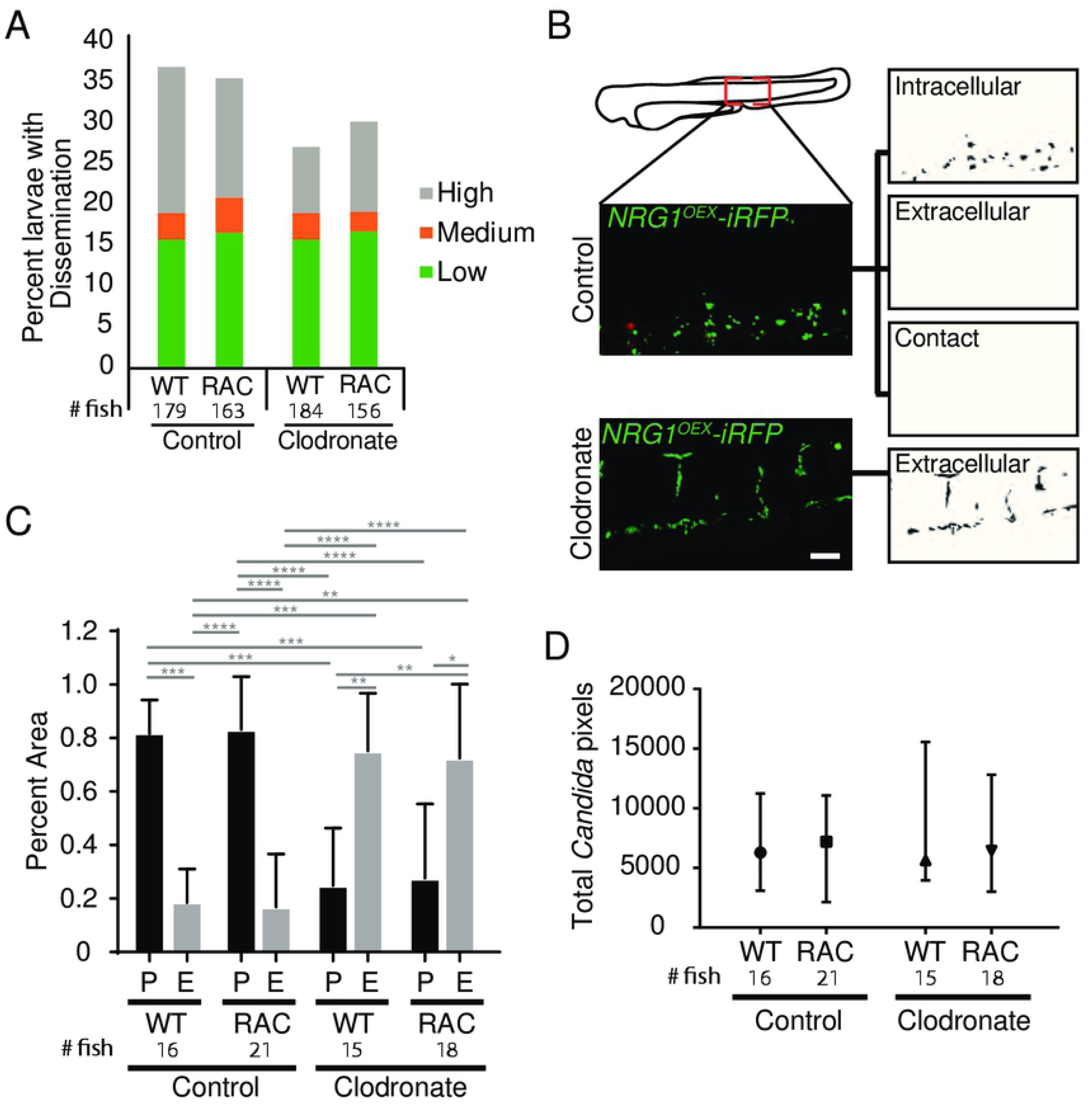
Phagocytes are not required for yeast dissemination to occur. Rac2-D57N and AB sibling larvae were injected with control or clodronate liposomes mixed with a 10 kDa dextran conjugated with Cascade Blue in the caudal vein at 28 hpf. Larvae were infected with the yeast locked *C. albicans* 4 hours later as described above. (A) Percent infected larvae with dissemination scored as “low” (1-10), “medium” (10-50), and “high” (>50 yeast) disseminated yeast at 40 hpi. Pooled from 6 experiments. Stats: Fisher’s Exact test, n.s. not significant p>0.05 (B) Method used to quantify disseminated yeast. Yeast were scored as Phagocytosed (P; inside or in close contact with a phagocyte) or Extracellular (E; not contained or in contact with phagocytes). Images were processed in ImageJ. Scale bar = 50 µm. (C) The proportion of all yeast either Phagocytosed or Extracellular for each treatment group. Stats: Kruskall-Wallis with Dunn’s posttest (* p≤ 0.05, ** p ≤ 0.01, *** p ≤ 0.001, **** p < 0.0001), and bars indicate the median with interquartile range. Pooled from 6 experiments. (D) Total pixel counts for quantified disseminated yeast for each group are shown with the median and interquartile range. Same larvae as in Panel C. Stats: Kruskall-Wallis with Dunn’s post-test. All comparisons were n.s.

### Phagocyte ablation is associated with increases in extracellular yeast in the bloodstream

We reasoned that elimination of both major professional immune cell types might alter infection dynamics significantly, limiting the ability of the host to contain bloodstream pathogens and thereby enhancing the ability of yeast to survive and proliferate once gaining access to the bloodstream. We quantified whether disseminated yeast were found intracellularly or extracellularly, as shown in Fig. 3B. While larvae without functional macrophages had a higher ratio of extracellular to intracellular yeast, neutrophil perturbation did not affect this ratio (Fig. 3C). Consistent with the categorical dissemination scores (Fig. 3A), the total number of disseminated yeast, as calculated in pixels, was not influenced by the absence of macrophages (Fig 3D). Together, these results suggest that although phagocytes, particularly macrophages, participate as Trojan Horses, and macrophages are required to contain disseminated yeast intracellularly, they are not required for dissemination to occur.

### TNFα production is dependent on macrophages but not neutrophils

These phagocyte ablation data suggested that neither phagocyte type is required for dissemination, but did not address if phagocyte ablation affects local inflammatory gene expression at the site of infection. To determine if *tnfa* expression was altered in response to macrophage ablation, *Tg(LysC:dsRed)/TgBAC(tnfa:GFP*) larvae with red neutrophils and TNF*a* promoter-driven GFP were infected and imaged for *tnfa* expression in presence or absence of macrophages. Representative images of control liposome- and clodronate liposome-treated larvae are shown in Fig. 4A, where the yellow outline depicts the infected area used for analysis. Neutrophils were found at the site of infection in both control and clodronate-treated larvae, but control larvae had slightly more neutrophil recruitment than their clodronate treated counterparts (Fig. 4B). TNFα expression was essentially absent in clodronate-treated infected larvae (Fig. 4C), correlating with a slight reduction in neutrophil recruitment to the infection site. We sought to determine if there was a relationship between TNFα and neutrophil levels in the clodronate-treated fish, which would indicate if neutrophils drive *tnfa* expression in the absence of macrophages. In control fish, we found a positive correlation between *tnfa:EGFP* expression and neutrophil levels, but in the absence of macrophages, this relationship was lost and there was even a slight negative correlation (Fig. S9A). Neutrophil recruitment to the site was only slightly reduced in larvae depleted of macrophages, but these larvae demonstrate near total loss of *tnfa:EGFP* expression (Fig. S9B). This is despite the fact that there is no significant difference in fungal burden among these cohorts (Fig. S9C). These experiments suggest that macrophages are the primary drivers of *tnfa* expression at the fungal infection site.

**Figure 4.**
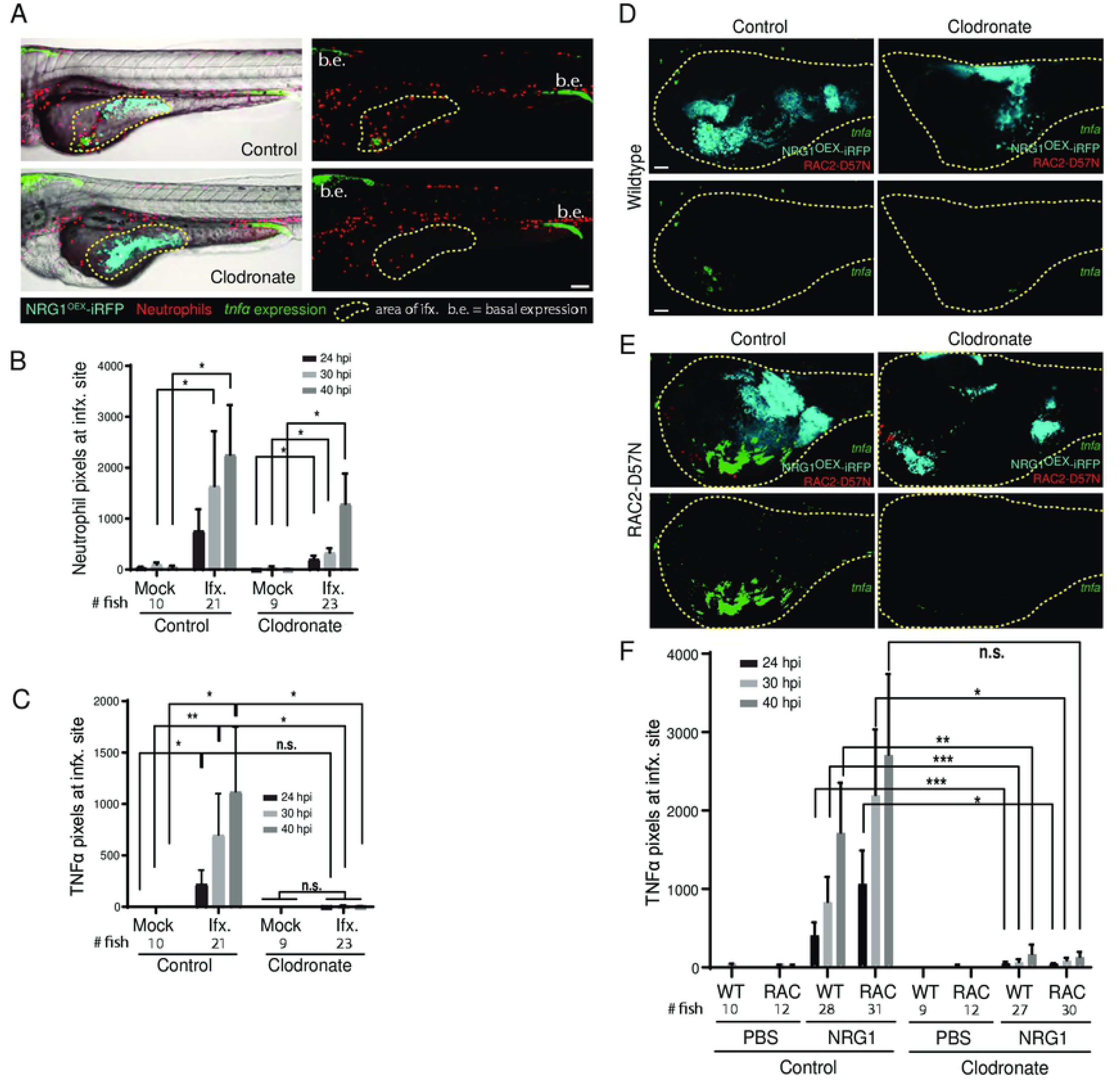
Macrophages are primarily responsible for TNFα production. *Tg(lysC:Ds-Red)/Tg(tnfα:GFP)* or Rac2-D57N/ *Tg(tnfα:GFP)* larvae with red fluorescent neutrophils and green fluorescence with *tnfα* expression were infected with a yeast-locked *C. albicans* as described. (A) Representative images of control and clodronate treated *Tg(LysC:Ds-Red)/Tg(tnfα:GFP)* larvae with functional neutrophils infected with a yeast-locked *C. albicans* at 40 hpi. Area in yellow outlines area of *Candida* growth in the yolk sac. Images were used to quantify neutrophil recruitment and *tnfα* expression. Scale bar = 100 µm. Markings are clarified in box below images. (B) and (C) Images were analyzed with ImageJ. (B) Neutrophil recruitment to the area of infection is not affected by macrophage ablation. Data pooled from 5 experiments, total fish used for quantification, left to right: n= 10, 21, 9, 23. Bars are the median with 95% confidence interval. Stats: Kruskall-Wallis with Dunn’s post-test. * p ≤ 0.05. (C) Larvae with ablated macrophages have an almost complete loss of *tnfα* expression. Bars are the means and SEM for ease of viewing. Stats: Mann-Whitney. * p ≤ 0.05, **p ≤ 0.01. (D) and (E) Representative images of control (left) and clodronate (right) treated Rac2-D57N x *Tg(tnfα:GFP)* sibling larvae infected with a yeast-locked *C. albicans* at 40 hpi. Area in magenta shows the yolk sac outline. Scale bar = 50 μm. (F) Larvae in which macrophages are ablated also have an almost complete loss of *tnfα* expression, which is independent of neutrophils. Pooled from 5 experiments. Stats: Kruskall-Wallis with Dunn’s post-test, * p ≤ 0.05, ** p ≤ 0.01, and bars are the mean and SEM.

To complement these macrophage experiments, we sought to examine the effects of macrophage ablation in the context of neutrophil inactivation. Rac2-D57N and *TgBAC(tnfa:GFP*) fish were crossed to generate larvae with functionally deficient neutrophils and TNFα-controlled green fluorescence. Representative images of control and clodronate-treated fish with median *tnfa:GFP* levels at 40 hpi suggest there is no *tnfa:GFP* expressed in macrophage ablated fish (Fig. 4D&E, Movies S6-9. Quantification of images of these fish demonstrate a trend toward higher *tnfa* expression in larvae with functional macrophages as compared to the clodronate-treated larvae, regardless of neutrophil status (Fig. 4F). Time-lapse movies further demonstrate macrophage-like cells turning on *tnfa* expression (Movie S10) and the lack of *tnfa* expression in macrophage-ablated fish (Movie S11). Similar to what was found previously (Fig. 3A), there was no change in dissemination rates with loss of one or both phagocytes types in our Rac2-D57N cross with *TgBAC(tnfa:GFP*) (data not shown). This data suggests that macrophages are vital for *tnfa* expression either because they are the primary producers of TNFα or because their presence is required so that other host cells are cued to produce TNFα.

### Loss of blood flow reduces overall number of disseminated yeast but does not affect dissemination frequency or tissue location

While many pathogens use phagocytes to traverse epithelial and endothelial barriers, regulated endo- and exocytosis and paracellular movement represent alternative pathways of yeast dissemination into the bloodstream [12, 28, 36]. Extracellular yeast that enter the bloodstream may be then transported away from the infection site by blood flow to distant tissues in the tail or head. To determine the role of blood flow in spreading infection, we took advantage of the ability of zebrafish larvae to grow and develop for the first seven days without a heartbeat, exchanging gases directly with the surrounding water [37–39].

We allowed for development until the time of infection to avoid disrupting normal hematopoiesis, then blocked blood flow using non-toxic doses of one of two previously characterized inhibitors of heart beat, terfenadine [40, 41] or valproic acid [42, 43]. Time-course and time-lapse imaging of treated larvae showed that neither drug quantitatively or qualitatively reduced phagocyte responses to *C. albicans* challenge, demonstrating that phagocyte recruitment is not dependent on blood flow (Fig. S10A-B; Movies S12 & S13; terfenadine- and valproic acid-treatment, respectively). Overall dissemination rates were also unchanged between larvae with and without blood flow in both *Tg(fli1:EGFP)* fish (Fig. 5A-B) and in *Tg(Mpeg:GAL4)/(UAS:nfsB-mCherry)/Tg(mpx:EGFP)* fish (Fig. S10C-D). However, we noticed a qualitative decrease in dissemination levels so we split fish into categories of different levels of dissemination. Interestingly, the proportion of fish with “High Dissemination” was lower with loss of blood flow (1% vs 10%, p<0.001; Fig. 5B) and a similar trend was seen in additional but underpowered experiments with fewer fish treated with either terfenadine or valproic acid (for terfenadine 0% vs. 9%, p=0.0686; for valproic acid 0% vs. 15%, p=1.000; Fig. 10C-D). This qualitative alteration in the pattern of dissemination suggested that blood flow is not required for spread but does play a limited role in spreading the infection.

**Figure 5.**
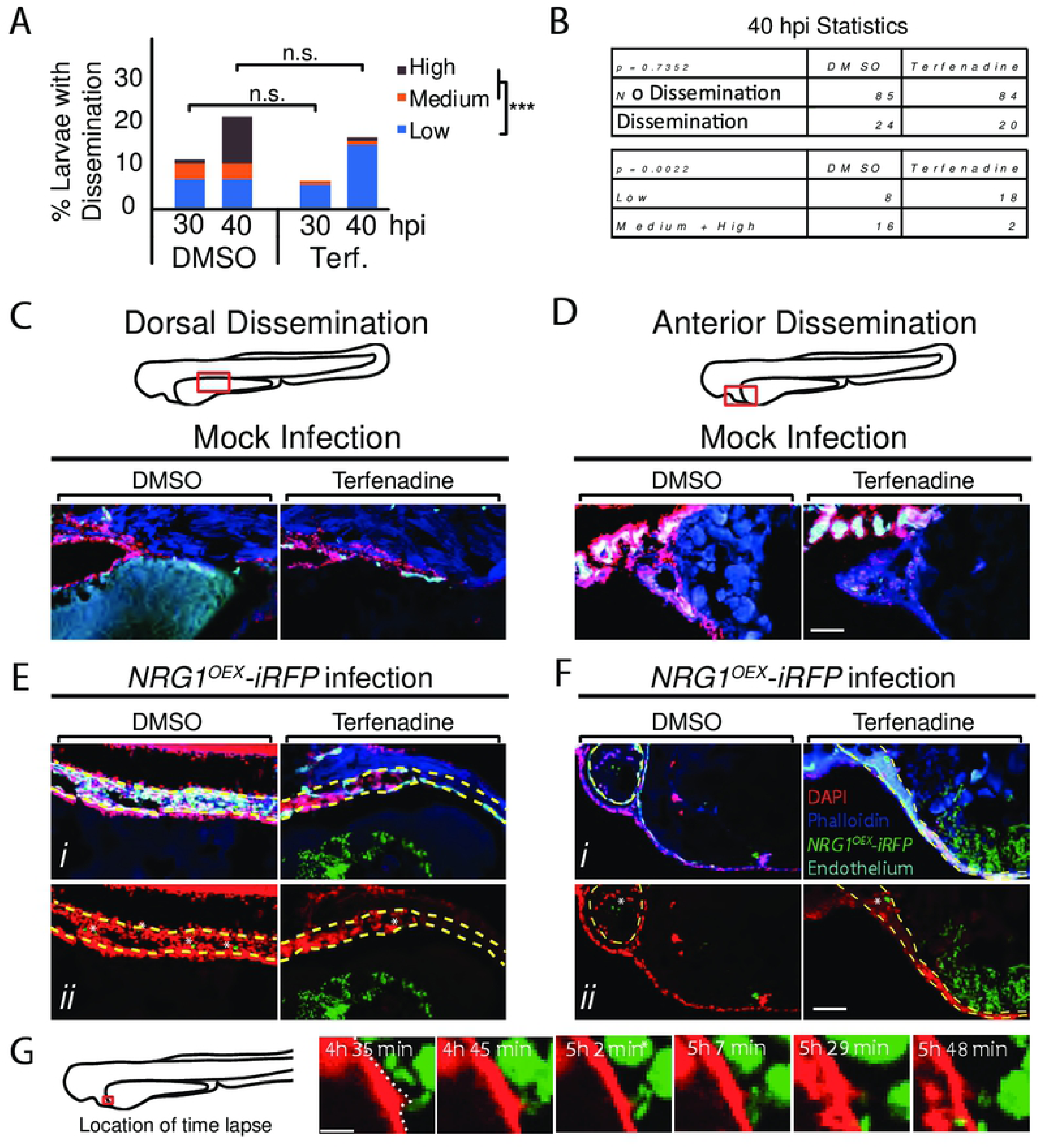
Blockade of blood flow partially limits dissemination. (A-G) *Tg(fli1:EGFP)* larvae, with GFP-expressing endothelial cells, were infected as previously described, treated with vehicle (DMSO) or terfenadine (2 μM) and processed for histology at 40 hpi. (A-B) Percent larvae with dissemination at 30 and 40 hpi scored as having “low” (1-10), “medium” (10-50), and “high” (>50 yeast) disseminated yeast. Pooled from 7 experiments, DMSO n = 110, Terfenadine n = 118. Fisher’s Exact test was used to test for differences between “low” and “medium” plus “high” scores at 40 hpi, *** p ≤ 0.001. (C-F) Sections of *Tg(fli1:EGFP)* mock infected (C-D) and *Candida*-infected (E-F) larvae compare dorsal dissemination (E) and anterior dissemination (F) events to the same tissue in mock-infected fish. Sections were stained with DAPI (nuclei, red) to indicate surrounding cells and phalloidin (actin; blue) to indicate host structures. NRG1^OEX^-iRFP (green) and *Tg(fli1:EGFP)* (cyan) retained fluorescence during sectioning and staining. White asterisks indicate disseminated yeast which appear to be embedded in the yolk syncytial layer and larval heart. Scale bar = 100 μm. (G) A vehicle-treated *Tg(fli1:EGFP)* (red) fish infected with NRG1^OEX^-iRFP (green) with intact blood flow was imaged by time-lapse. Area in red box is the focus of the time-lapse, with frames taken from Movie S14 a cropped area of the complete time lapse included as Movie S15 Scalebar = 10 μm.

To explore further whether blockade of blood flow alters the qualitative nature of yeast dissemination, we sought to characterize the context of fungal spread into the fish at higher spatial and temporal resolution. To enhance spatial resolution and assess the entire tissue context of disseminating yeast, we performed confocal microscopy on frozen sections rather than solely focusing on fluorescently marked cells. Examination of mock-infected fish confirmed that drug treatment caused no overall loss of tissue structure either dorsal or anterior to the yolk (Fig. 5C-D). Histology of infected fish revealed that yeast that have left the yolk are often found in tissues near the thin endothelial layer surrounding the yolk in both vehicle- and terfenadine-treated larvae with blocked blood flow, either dorsally (Fig. 5E) or anteriorly (Fig. 5F) of the yolk. The similar tissue locations of yeast with and without blockade of circulation suggest that the dissemination path of yeast near the yolk is not affected by eliminating blood flow. Further, these high-resolution images show that yeast are clearly found within blood vessels with endothelial cells expressing the *fli1:EGFP* reporter, pseudocolored in cyan [44]. We also were able to observe the dynamic movement of yeast from the yolk to the blood vessel through time-lapse imaging (Fig. 5G, Movie S14. The frames in Fig. 5G demonstrate the movement of yeast through the endothelial cell layer directly from the yolk sac in a vehicle-treated larva. While the movie clearly shows movement into the blood vessel and adherence in the context of strong blood flow, we are unfortunately unable to deduce whether this was host micropinocytosis or paracellular invasion. Nevertheless, given that the movement occurs through an endothelial barrier that appears intact after the migration event, this time lapse suggests how host endothelial cells and/or phagocytes may play a role in the passage of yeast from a localized infection site to the bloodstream.

### Spread of yeast to distant tissues is inhibited with loss of phagocyte activity and blood flow

Blockade of blood flow reduces the number of fungi that leaving the infection area but does not affect the percent of fish with some tissue-to-blood dissemination. Because phagocyte responses (Fig. S10) were not reduced or eliminated by blocking heartbeat, we reasoned that phagocytes might account for basal rates of dissemination. To test the combined activity of blood flow and phagocyte-mediated spread, we quantified dissemination in larvae lacking phagocyte activity and heartbeat using *mpx:Rac2-D57N* to block neutrophil function, clodronate liposomes to ablate macrophages and terfenadine to block blood flow. As expected, based on our previous experiments (Fig. 4), larvae lose *tnfα* expression with macrophage ablation, which is not correlated with the amount of *Candida* in the yolk (Fig. S11). Rac2-D57N larvae treated with clodronate and terfenadine were found to have disseminated yeast limited to tissues adjacent to the yolk, while disseminated yeast in control larvae were transported to more distant places (Fig. 6A). As qualitatively scored based on the number of *C. albicans* outside of the yolk, loss of blood flow did not affect the overall percentages of fish with dissemination (Fig. 6B). To test whether blood flow moved disseminated yeast far from the infection site, we then quantified this disparity in near (<25 pixels from yolk) versus far (>55 pixels from yolk) dissemination (Fig. 6C). Strikingly, there was a significant difference in the pattern of yeast dissemination when macrophages were ablated and blood flow was blocked, with a reduction in distant dissemination (Fig. 6D). Statistically, there were no significant differences in the number of yeast outside but close to the yolk in any condition (Fig. 6E). In contrast, loss of blood flow in the context of macrophage ablation caused a significant decrease in the number of distantly-disseminated yeast (Fig. 6F). This loss of extensive dissemination was even more severe when neutrophils were also inactivated. Together, these data suggest that a combination of host activities play important roles in the spread of yeast from the infection site.

**Figure 6.**
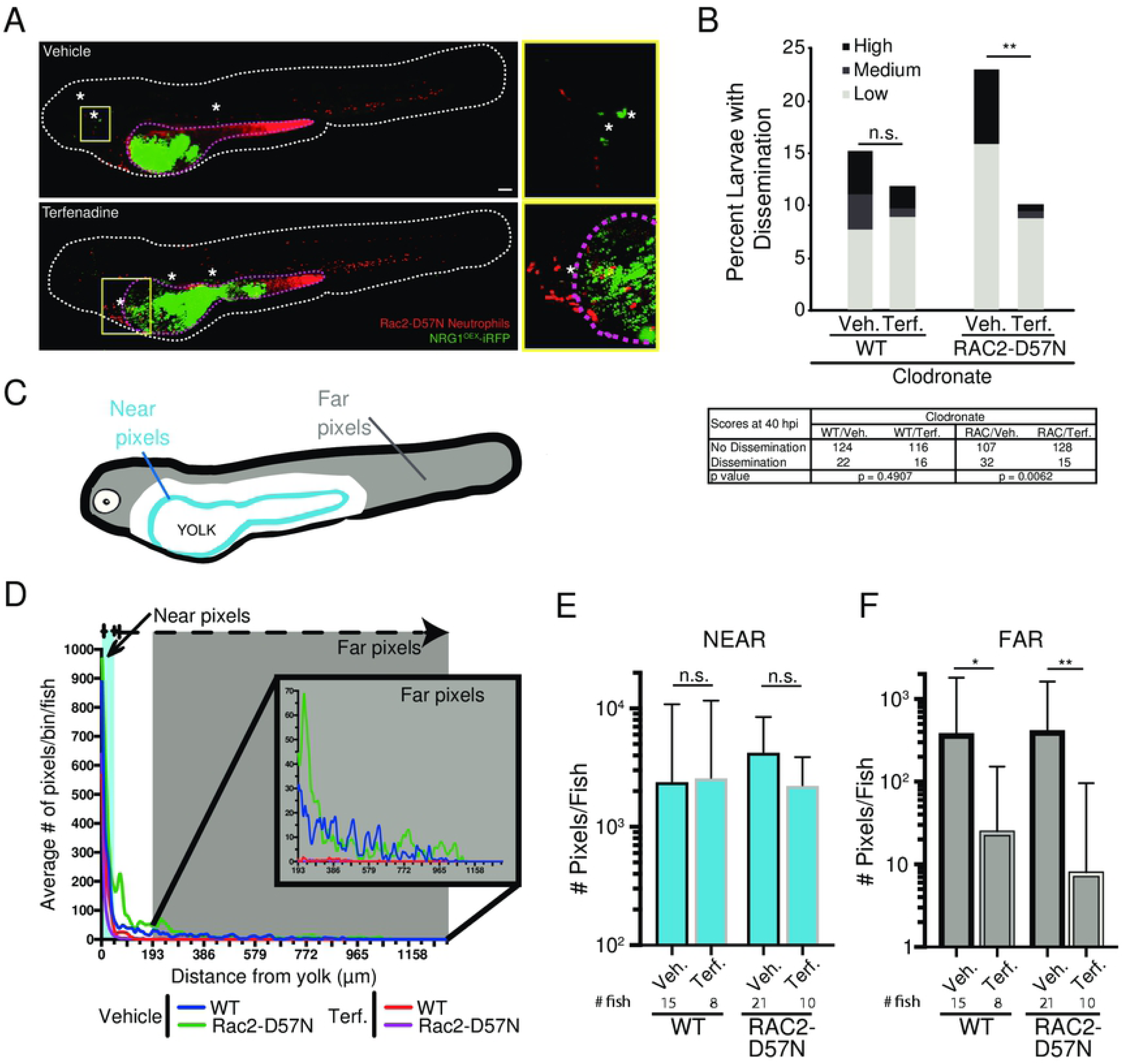
Dissemination is inhibited with loss of phagocytes and blood flow. Rac2-D57N zebrafish were crossed with AB fish for neutrophil-deficient or wild type offspring. All larvae were injected with clodronate liposomes for macrophage ablation and bathed in vehicle (DMSO) or 2 μM terfenadine for blood flow blockade. Larvae were then infected with NRG1^OEX^-iRFP. (A) Representative Rac2-D57N larvae with dissemination either with (Vehicle) or without (Terfenadine) blood flow. The white line outlines the fish body, the magenta line indicates the yolk sac, and white asterisks denote yeast that are disseminated. The yellow box indicates an area with disseminated yeast that has been magnified at the right. Scale bar = 100 µm. (B) Percent larvae with scores of dissemination at 40 hpi, pooled from 6 experiments. Stats: Fisher’s exact test, * p ≤ 0.05. (C) Schematic of the scoring system for disseminated yeast. Yeast that were ≤25 pixels from the yolk sac edge were scored as “near” and yeast that were ≥55 pixels from the yolk sac edge were scored as “far”. (D) A frequency distribution histogram of the distance in pixels that yeast travel from the yolk sac edge sorted into single bins. Distances less than 5 pixels was omitted from analysis. Same fish quantified as in C and D. Pooled data from 54 larvae. (E) Total number of near yeast pixels per fish, shown as median and confidence interval (F) Total number of far yeast pixels per fish, shown as median and confidence interval. (D-F) Pooled from 6 experiments. Stats: Mann-Whitney, n.s. not significant, * p ≤ 0.05, ** p ≤ 0.01

### Dissemination of wildtype yeast follows similar pattern as yeast-locked yeast

These data, taken together, suggest that yeast spread to distant tissues by a combination of phagocyte-mediated dissemination and extracellular dispersal through the epithelial and endothelial layers followed by circulation-mediated movement. While use of the yeast-locked NRG1^OEX^ strain enabled these detailed analyses of dissemination in the absence of extensive tissue damage and early death, we felt it was important to determine if key aspects of yeast dissemination are mirrored during infections with wildtype *C. albicans* that produce a mix of filaments and yeast. Infections with wildtype *C. albicans*—at a low temperature that promoted production of both yeast and filaments—resulted in the expected high levels of mortality due to invasive filaments [5]. Nevertheless, we were able to confirm that phagocyte recruitment precedes dissemination (Fig 7A and B), and dissemination frequency is unaffected by loss of phagocyte function (Fig. 7C). Furthermore, phagocyte recruitment to the site of infection is accompanied by a macrophage-dependent local upregulation of TNFα when neutrophils are disabled (Rac2-D57N) but not when neutrophils are functional (Fig. 7D and E; Movies S15-16. These contributions by neutrophils may reflect neutrophil-hyphal interactions present only with the wildtype *C. albicans*. Many interactions of macrophages and neutrophils were captured in these experiments (Movies S15-S17). Consistent with previous work that showed no association of fungal burden with dissemination [5], we found no differences in fungal burden either between fish with and without dissemination or between immunocompetent versus immunocompromised fish (data not shown). Taken together, these data suggest that the processes of yeast dissemination are similar for both yeast-locked or wildtype yeast.

**Figure 7.**
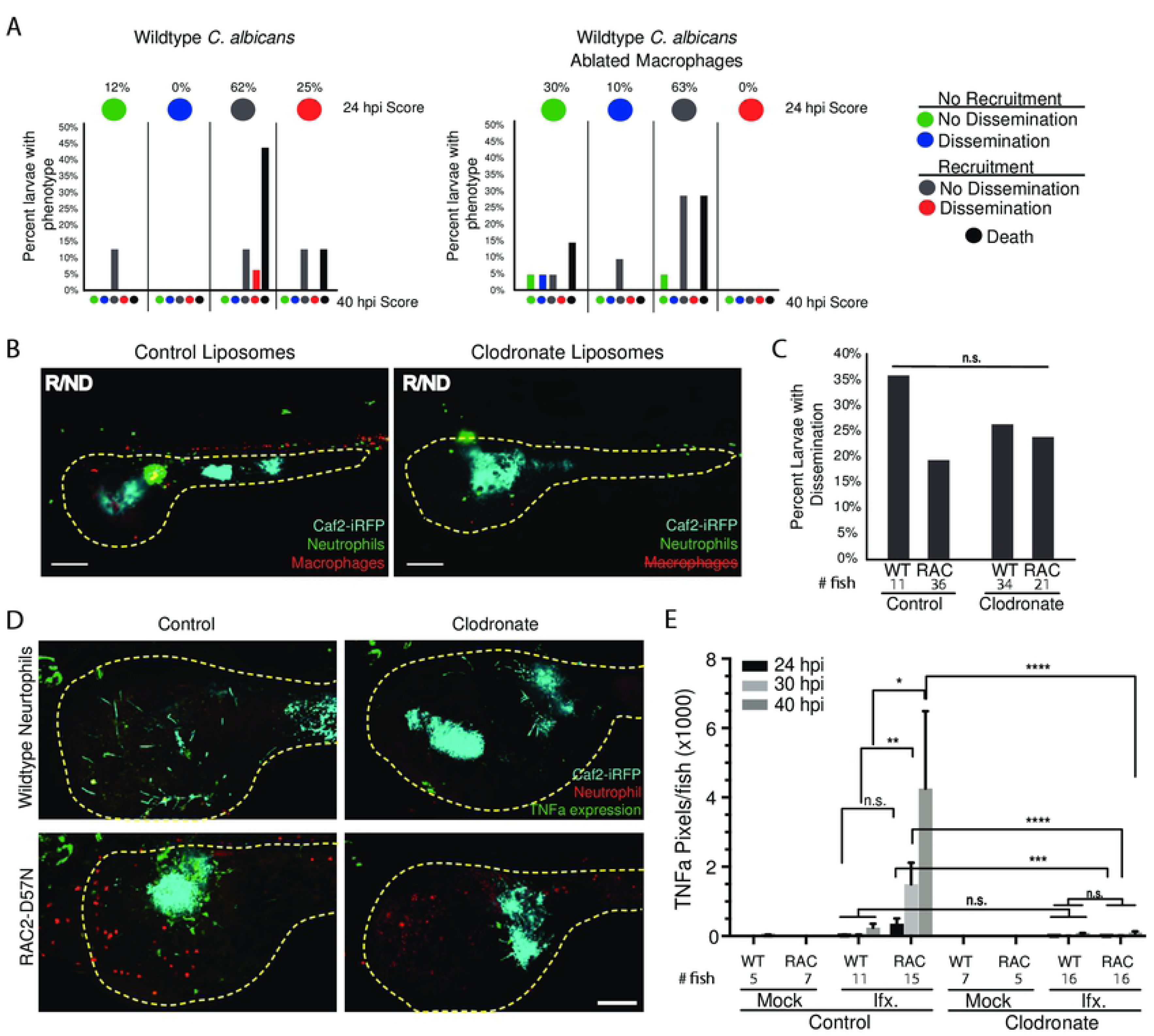
Wild type *C. albicans* mirrors yeast-locked strain. *Tg(Mpeg:GAL4)/(UAS:nfsB-mCherry)/Tg(mpx:EGFP)* or RAC2-D57N/*Tg(tnfα:GFP)* larvae were injected with control or clodronate liposomes and infected with the wild type Caf2-iRFP *C. albicans* as described for the yeast-locked infections. (A) Dissemination dynamics for wildtype fungi are similar to dynamics for yeast-locked and are unaffected by macrophage ablation. *Tg(Mpeg:GAL4)/(UAS:nfsB-mCherry)/Tg(mpx:EGFP)* were infected and scored for immune recruitment and dissemination of yeast between 24 and 40 hpi, as described in Fig. 1B. Top circles represent the percentages of fish at 24 hpi, colored with the indicated recruitment/dissemination phenotype. Bottom bars represent phenotypes at 40 hpi, grouped based on their 24 hpi phenotype. Pooled from 2 experiments, control larvae n = 16, clodronate larvae n = 21. (B) Representative images of Recruited/Non-disseminated *Tg(Mpeg:GAL4)/(UAS:nfsB-mCherry)/Tg(mpx:EGFP)* larvae treated with control or clodronate liposomes at 40 hpi. Area in yellow shows the yolk sac outline. Scale bar = 50 μm. (C) Dissemination of wildtype yeast does not require intact phagocytes. Percent dissemination of *Tg(Mpeg:GAL4)/(UAS:nfsB-mCherry)/Tg(mpx:EGFP)* larvae (left) and RAC2-D57N/*Tg(tnfα:GFP)* larvae (right) at 40 hpi. Right: Pooled from 2 experiments as in panel A. Left: Pooled from 3 experiments. Fisher’s exact test, n.s. (D) Wildtype elicits a macrophage-dependent expression of TNFα. Representative images of control (left) and clodronate (right) treated Rac2-D57N/*Tg(tnfα:GFP)* larvae. Magenta outlines the yolk sac. Scale bar = 100 μm. (E) Pixel area of cells expressing *tnfα* during wild type *C. albicans* infection. Pooled from 3 experiments. Stats: Kruskal-Wallis with Dunn’s post-test, * p ≤ 0.05, ** p ≤ 0.01, *** p ≤ 0.001., **** p ≤ 0.0001. Bars indicate the mean and SEM.

## Discussion

Although invasive candidiasis poses a significant clinical risk, we still understand little about how this small non-motile fungus spreads throughout a host and what roles the host itself plays in limiting or enabling its movement. Using longitudinal intravital imaging of *C. albicans* and the host, we found that phagocyte-dependent and -independent mechanisms provide redundant pathways from tissue to bloodstream and throughout the host. On one hand, we find phagocytes can traffic yeast far from the infection site even in the absence of blood flow, releasing them in distant tissues. On the other hand, we show that even when phagocytes are disabled the yeast are able to efficiently get into the bloodstream and use blood flow to reach far tissues. These multiple strategies emphasize the versatility of *C. albicans* and suggest that we need to understand fungal interactions with both endothelial cells and phagocytes to understand the mechanics of tissue-to-bloodstream dissemination.

Phagocyte recruitment correlates with pro-inflammatory gene upregulation and local *tnfα* expression. Cytokine upregulation is a key step in innate immune response to an infectious threat, and this is the case in our infection model [45]. Imaging by time-lapse at the single cell level revealed that *tnfα*-expressing cells are motile and are absent when macrophages are depleted, suggesting that only macrophages express *tnfα* in this infection model. Although imaging of *tnfα* levels in macrophage transgenics was not performed here, nearly all *tnfα*-GFP+ cells were found to also be *mpeg1:mCherry*+ in a similar swimbladder infection model [46]. Proinflammatory cytokines at the infection site likely promote vascular permeability that may enhance fungal spread to the bloodstream by endocytic, paracellular and/or Trojan-horse pathways [7, 22, 23, 47].

We documented all of the key steps of Trojan Horse-mediated spread of *C. albicans in vivo*: recruitment, phagocytosis, reverse migration and fungal escape. The hijacking of phagocytes by *C. albicans* raises important questions about how and why innate immune cells transport yeast into the bloodstream and release them far from the infection site. While neutrophil reverse migration is now well-documented [48–52], we still do not know what host and fungal signals regulate yeast-laden macrophages to leave the infection site. Further, although non-lytic exocytosis (NLE) is utilized by many pathogens, its regulation is poorly understood and likely involves both host and pathogen cues [24, 53]. Further work focused on NLE *in vivo* may reveal more about how *C. albicans* and other pathogens escape macrophage containment during vertebrate infection.

Neutrophils and macrophages were found to be dispensable for efficient dissemination. This was unexpected because we documented Trojan Horse dissemination, we found that phagocytes are sufficient to spread *C. albicans* in the absence of blood flow, and we know that macrophages are key for spread of other fungal and bacterial pathogens [24, 53]. *C. albicans* may be able to efficiently spread without hitchhiking on phagocytes because it grows relatively well extracellularly compared to other phagocyte hitchhikers and/or because it has effective alternative means of entering and exiting the bloodstream without phagocytes. Further, as phagocytes both limit infection and spread yeast, there may be a counterbalancing effect of phagocyte elimination, where extracellular yeast are able to survive and proliferate in the blood in their absence. In this scenario, a decrease in phagocyte-mediated spread could be made up for by an increase in survival and proliferation in the bloodstream in the absence of attacking immune cells.

How do *C. albicans* yeast get through the epithelium and endothelium without phagocytes? Hyphae are known to translocate through epithelial and endothelial cell barriers *in vitro*, but yeast translocation has not yet been analyzed in these systems. Yeast passage through epithelial layers is infrequent in most epithelial *in vitro* models of barrier passage, which could be due to the loss of normal tissue architecture *in vitro* and/or to the altered expression of ligands and receptors that mediate internalization. There are a number of candidate host receptors, as well as *C. albicans* adhesins and proteolytic enzymes that could participate in yeast translocation, including cadherins, EphA2, EGFR, ErbB2, secreted aspartyl proteases (SAPs), candidalysin, Als3p [7, 54, 55].

Blood flow and phagocyte-mediated dissemination represent redundant mechanisms for *C. albicans* to spread throughout the host body from a localized infection. While it had been postulated that both strategies might be used, full-animal imaging of infection progression enabled the quantification of both mechanisms at an unprecedented level. Interestingly, loss of blood flow alone reduces the amount of disseminated yeast within fish with dissemination but not rates of dissemination into the bloodstream or far from the infection site. On the other hand, elimination of phagocyte function alone affects neither overall levels of spread nor frequency. The dissemination of yeasts adjacent to the yolk in the absence of heart beat and phagocyte function suggests that diffusion through blood vessels isn’t sufficient to carry away yeast, perhaps because of adherence to the vascular lumen [56, 57]. It is intriguing that the two mechanisms are not additive, highlighting the potentially double-edged sword of phagocytes that can enhance dissemination through Trojan horse-mediated mechanisms but also limit spread of free yeast through phagocytosis and containment. The higher level of dissemination and greater amount of extracellular yeast we observe in phagocyte-incapacitated hosts (Fig. 3 and Fig. S7) is consistent with an important role for phagocytes in limiting fungal proliferation in the bloodstream. Tracking the fate of individual phagocytes by adapting the Zebrabow fish line [58, 59] and/or imaging individually photoswitched yeast should allow single-cell quantification of these activities in the future.

While the larval zebrafish model has unique advantages in imaging and manipulation, these come with some limitations as well. The small size of the fish allows one to image the whole fish on the confocal, but the related limitation of the model is that dissemination distances are quite small, compared to what you would have in a mouse or human. Therefore, in translating our findings to the human host, we may find a differential time frame for extracellular spread by rapid blood flow as compared to phagocytes moving more slowly through tissue and lymph. Furthermore, although the overall anatomy is similar between fish and man, the tissue architecture of yolk, epithelium and endothelium is not representative of all potential sites of tissue-to-bloodstream dissemination in mammals. Based on our findings in the zebrafish and the known conservation of cell types and molecules among vertebrates, we predict that both phagocyte-mediated and extracellular mechanisms of spread are important in mammalian tissue-to-blood dissemination. Unfortunately, although large groups of fungi can be tracked non-invasively in the mouse, testing of these ideas in a mammal will have to wait for new methods that will allow tracking of individual yeast and phagocytes to distant tissues [60–67].

Leveraging the advantages of the zebrafish model system, we have quantified two redundant means of *C. albicans* yeast spread from tissue to blood. We documented Trojan Horse spread of *C. albicans* and showed that it is redundant with phagocyte-independent dissemination. Conservation of adhesion molecules, cell types and anatomy among all vertebrates suggests that both mechanisms are likely to also be important in mammalian infection [68–71]. Clinical observations are consistent with our findings that *C. albicans* moves in the blood both inside and outside of phagocytes [72–75]. From a therapeutic standpoint, our results suggest that prevention of *C. albicans* dissemination will require interventions that block both endothelial and phagocyte-driven movement of yeast.

## Materials and Methods

### Animal care and maintenance

Adult zebrafish used for breeding were housed at the University of Maine Zebrafish Facility in recirculating systems (Aquatic Habitats, Apopka, FL). All zebrafish studies were carried out in accordance with the recommendations in the Guide for the Care and Use of Laboratory Animals of the National Research Council [76]. All animals were treated in a humane manner and euthanized with Tricaine overdose according to guidelines of the University of Maine Institutional Animal Care and Use Committee (IACUC) as detailed in protocols A2015-11-03 and A2018-10-01. Following collection, embryos were kept 150 mm petri dishes with E3 media (5 mM sodium chloride, 0.174 mM potassium chloride, 0.33 mM calcium chloride, 0.332 mM magnesium sulfate, 2 mM HEPES in Nanopure water, pH 7) plus 0.3 mg/liter methylene blue (VWR, Radnor, PA) to prevent microbial growth for the first 6 hours. Larvae were then moved to fresh E3 media supplemented with 0.02 mg/ml of 1-phenyl-2-thiourea (PTU) (Sigma-Aldrich, St. Louis, MO) to prevent pigmentation. Larvae were reared at 28°C at a density of 150 larvae per 150 mm petri dish. Transgenic lines used are provided in Table 1.

**Table 1.** Zebrafish lines used

| **Transgenic line** | **Reference** |
| --- | --- |
| *Tg(mpx:EGFP)i11rTg* | [77] |
| *Tg(mpeg1:GAL4)gl24Tg* | [34, 78] |
| *Tg(UAS-E1b:NTR-mCherry)c264Tg* | [79] |
| *Tg(Mpeg1:GAL4)/(UAS:Kaede)* | [32] |
| *TgBAC(tnfa:GFP)pd1028Tg* | [22] |
| *Tg(LysC:dsRed)* | [22, 80] |
| *Tg(fli1:EGFP)* | [44] |
| *Tg(mpx:mCherry,rac2_D57N)zf307Tg* | [35, 81] |
| AB (wild type) | Zebrafish International Resource Center |

**Table 2.** Candida albicans strains used

| *C. albicans* Strain | Parent Strain | Figure | Genotype | Reference |
| --- | --- | --- | --- | --- |
| NRG1<sup>OEX</sup>-iRFP | THE21 | All other experiments | *ade2::hisG/ade2::hisG*<br>*ura3::imm434/ura3::imm434::URA3- tetO ENO1/eno1::ENO1 tetR- ScHAP4AD-3XHA-ADE2 pENO1-iRFP-NATR* | This study, [5, 13] |
| *efg1Δ/Δ cph1Δ/Δ*-iRFP | CAI4 | Fig S8 | *ura3::1 imm434/ura3::imm434*<br>*cph1::hisG/cph1::hisG*<br>*efg1::hisG/efg1::hisG-URA3-hisG*<br>*pENO1-iRFP-NATR* | [5, 82] |
| *efg1Δ/Δ cph1Δ/Δ*-dTomato | CAI4 | Movie S5 | *ura3::1 imm434/ura3::imm434*<br>*cph1::hisG/cph1::hisG*<br>*efg1::hisG/efg1::hisG-URA3-hisG*<br>*pENO1-dTomato-NATR* | [5, 82] |
| NRG1<sup>OEX</sup>-dTomato | THE21 | Fig 1B | *ade2::hisG/ade2::hisG*<br>*ura3::imm434/ura3::imm434::URA3- tetO ENO1/eno1::ENO1 tetR- ScHAP4AD-3XHA-ADE2 pENO1-dTomato-NATR* | [5, 82] |
| Caf2-dTomato | CAF2.1 | Fig S3 | *Δura3::imm434/ URA3* pENO1-dTomato-NATR | [84] |
| Caf2-iRFP | CAF2.1 | Fig 4, Movies S15-17 | *Δura3::imm434/ URA3* pENO1-iRFP-NATR | [83] |

**Table 3.**
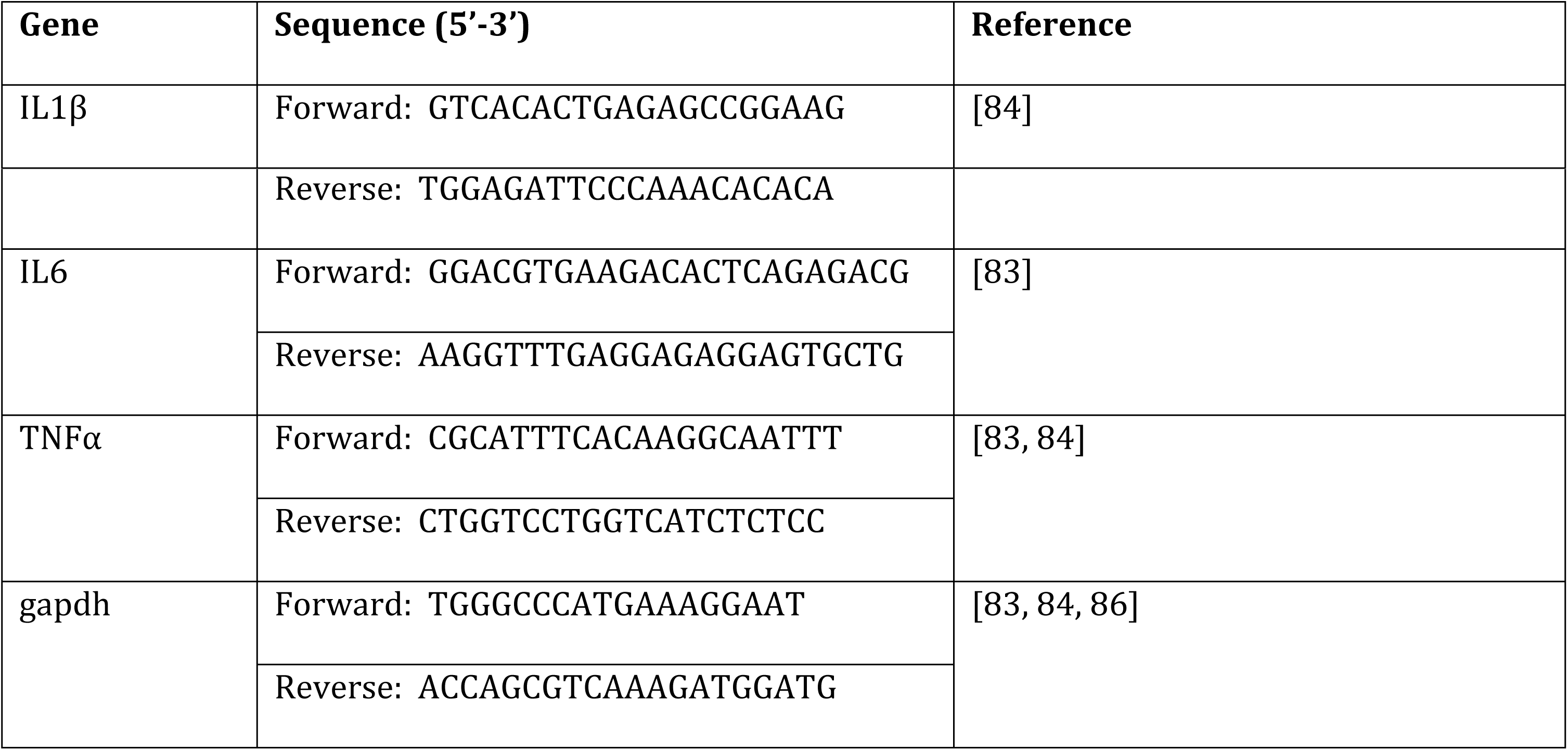
qPCR Primers

### Ethics statement

All zebrafish studies were carried out in accordance with the recommendations in the Guide for the Care and Use of Laboratory Animals of the National Research Council [76]. All animals were treated in a humane manner and euthanized with Tricaine overdose according to guidelines of the University of Maine Institutional Animal Care and Use Committee (IACUC) as detailed in protocols A2015-11-03 and A2018-10-01.

### Zebrafish larvae infection

Larvae at ∼32 hours post fertilization were manually dechorionated and anesthetized in fresh E3 media plus 0.02 mg/ml PTU and Tris-buffered tricaine methane sulfonate (160 μg/ml; Tricaine; Western Chemicals, Inc., Ferndale, WA). Larvae were injected in the yolk sac with 5 nl volume PBS control or *C. albicans* at 5×10^6^ CFU/ml in PBS. Larvae were microinjected in the yolk as described previously [5] and screened on a Zeiss Axio Observer Z1 microscope to ensure the correct injection and starting inoculum of *C. albicans* in the yolk (Carl Zeiss Microimaging, Thornwood, NJ). Scored larvae were then moved to fresh E3 media plus PTU which was changed every other day. Where indicated, media was supplemented with metronidazole, terfenadine, or valproic acid following infection. Larvae were kept at 28°C for NRG1^OEX^ yeast locked infections [82] or 21°C for wild type Caf2 infections [5]. These temperatures were chosen as they are safest for zebrafish and allow for yeast and hyphal growth of *C. albicans* [5, 78]. Fish were scored for levels of dissemination using the scale: 1-10 yeast (low), 11-50 (medium), >50 (high). Following 42 hours post infection, larvae were euthanized by Tricaine overdose. Mortality was observed as loss of heartbeat in all experiments where bloodflow was not altered. In experiments utilizing valproic acid or terfenadine to reduce heart rate and blood flow, mortality was scored as larval putrefaction (loss of tissue integrity, graying of tissue, and sloughing of dermis layer).

### *C. albicans* strains and growth conditions

*C. albicans* strains used for larval infection were grown for 24 hours at 37°C on yeast-peptone-dextrose (YPD) agar (20 g/L glucose, 20 g/L peptone, 10 g/L yeast extract, 20 g/L agar, Difco, Livonia, MI). Single colonies were picked to 2 x 5 mL liquid YPD and grown overnight at 30°C on a wheel. Prior to microinjection, liquid cultures were washed twice in phosphate buffered saline (5 mM sodium chloride, 0.174 mM potassium chloride, 0.33 mM calcium chloride, 0.332 mM magnesium sulfate, 2 mM HEPES in Nanopure water, pH = 7) and the concentration of yeast adjusted to 5×10^6^ CFU/ml in PBS for larval zebrafish infections. Wild type Caf2-dTomato *C. albicans* was used for heat killing and UV inactivating experiments. Overnight cultures were washed twice and resuspended in PBS at 2.5×10^7^ cells/ml, then boiled for 10 minutes or placed in an uncovered polystyrene petri dish (60 mm x 15 mm, VWR, Radnor, PA) for exposure. A CL-1000 UV cross-linker was used for UV inactivation of *C. albicans* [83] (UVP, Vernon Hills, IL) and yeast were exposed four times to 100,000 μJ/cm^2^ with swirling between each exposure. Following boiling or UV-inactivation, cells were stained with AlexaFluor 555 by coincubation of cells in PBS with sodium bicarbonate (0.037 M final concentration, pH 8.2) in the dark for 40 minutes with periodic vortexing. Cultures were then washed four times in PBS and brought to 2×10^7^ CFU/ml for injection into the yolk sac. Proper heat killing and UV inactivation was confirmed by plating 50 μl of the prepared 2×10^7^ CFU/ml killed yeast and 50 μl of the prepared 5×10^6^ CFU/ml live yeast used for larval injection on YPD plates and incubated overnight at 30°C.

For CFU quantification, groups of 6 larvae were taken after injection of *C. albicans* and screening and homogenized at 0 hpi in 600 ul of PBS. 100 μl of the homogenate was plated on YPD agar supplemented with gentamicin (30 μg/ml, BioWhittaker, Lonza), penicillin-streptomycin (250 μg/ml, Lonza), and vancomycin hydrochloride (3 μg/ml, Amresco, Solon, OH). Individual larvae were homogenized for CFUs in 600 μl PBS after the 40 hpi timepoint, and 100 μl of the 1:6 dilution or 1:60 dilution was plated on YPD for countable colonies. Plates were labeled by plate wells so that the larvae CFU count could be paired with dissemination scores and larval images. YPD plates were incubated overnight at 30°C and colonies counted the following day so CFUs/fish could be calculated.

### Macrophage Ablation

Where indicated, macrophages were ablated by injection of clodronate liposomes (Clodronate liposomes, Amsterdam, The Netherlands) in the caudal vein, larvae were bathed in metronidazole, or embryos were injected with *pu.1* morpholino oligonucleotides (MO, Genetools). Liposomes were injected in the caudal vein at a 3:1:1:1 ratio of 5 mg/ml liposome stock:PBS:phenol red:10 kDa Cascade blue dextran (8% w/v) in a total volume of 8-10 nl per larvae [9] at ∼28 hpf. Larvae recovered in E3 media for 4 hours and were then infected with *C. albicans*. *Tg(Mpeg:GAL4)/(UAS:nfsB-mCherry)* larvae were bathed in 20 mM metronidazole for 4 hours after injection with *C. albicans* yeast and then kept in E3 media plus PTU and 10 mM metronidazole for the remainder of the experiment [34, 78, 85]. Translational blocking (CCTCCATTCTGTACGGATGCAGCAT) and splice blocking (GGTCTTTCTCCTTACCATGCTCTCC) MOs were co-injected into 1-2 cell stage embryos [10, 30]. The MOs were combined (2.5 ng translation blocking and 0.4 ng splice blocking) with 10 kDa Cascade blue dextran (8% w/v) and phenol red for visualization of injection with total volume injected at 2 nl. Ablation of macrophages with MO technology was observed by loss of red fluorescence in *Tg(Mpeg:GAL4)/(UAS:nfsB-mCherry)* larvae.

### Chemical blockade of blood flow

For chemical inhibition of blood flow, larvae were reared and infected with *C. albicans* as described and then placed in petri dishes containing 0.1 mg/ml valproic acid in E3 media plus PTU or 2 μM terfenadine in E3 media plus PTU and corresponding vehicle controls (water and DMSO, respectively). Larvae were observed for reduced heartbeat and loss of blood flow in the trunk and tail by 24 hours post treatment (or 24 hpi). An Olympus SZ61 stereomicroscope system (Olympus, Waltham, MA) connected to an Excelis HDS microscopy camera and monitor system (World Precision Instruments, Sarasota, FL) was used to record 60 second movies with CaptaVision software. Movies were adjusted for frame rate and brightness in ImageJ-Fiji.

### RNA isolation and qPCR analysis

Larvae were pooled into groups based on phagocyte recruitment to the yolk sac and dissemination of yeast (non-recruited/non-disseminated, non-recruited/disseminated, recruited/non-disseminated, and recruited/disseminated). Larvae were euthanized by overdose in tricaine and immediately homogenized in TRIzol (Invitrogen, Carlsbad, CA) for RNA isolation. Larval groups ranged from 2-15 fish in 3 independent experiments. RNA was isolated using the Direct-zol RNA miniprep kit (Zymo Research, Irvine, CA) using the manufacturers recommended protocol. Final RNA was eluted in 20 ul nuclease free water and stored at −80°C until cDNA synthesis. Using the iScript reverse transcriptase supermix (Bio-Rad, Hercules, CA), cDNA was synthesized from 500 ng RNA per sample. RT-qPCR was done on a CFX96 thermocycler (Bio-Rad, Hercules, CA) using the cycles: 95°C for 30 s, 95°C for 5 s followed by 60°C for 20 s for 39 cycles, then 95°C for 10 s followed by 65°C for 5 s [83]. The Bio-Rad CFX Manager software was used to analyze threshold cycles and dissociation curves. Larval gene expression was normalized to the *gapdh* control gene (ΔCT), as done previously [83, 84], and compared to the mock infected PBS controls (ΔΔCT).

### Histology

Larvae were euthanized after 42 hpi in tricaine overdose and shipped overnight in PBS. Larvae were then fixed in 4% PFA at 4°C overnight, washed in PBS, and placed in 30% sucrose (MP Biomedicals 821713) in PBS for overnight or until the samples sank. Larvae samples were embedded in Tissue-Tek O.C.T. compound (OCT; Sakura Finetek 4583) and were kept at −80°C for at least 10 mins before sectioning. Sagittal 20-micron sections were collected with a Leica cryostat (CM3050S) and the sections were stained with DAPI (1 mg/ml; 1:2000 dilution) and Alexa Fluor™ 568 Phalloidin (Invitrogen A12380; 1:500 dilution) for 30 mins – 1 hour at room temperature. Then the slides were washed with PBS and mounted for imaging. Slides were then shipped back and sections were imaged on the Olympus Fluoview 1000 confocal microscope. Images were processed and pseudocolored in ImageJ.

### Fluorescence microscopy

*Tg(mpeg1:GAL4)* adults were crossed with *Tg(UAS:Kaede)* (yeast locked experiments) for larvae with photoconvertible green to red macrophages [78]. Larvae were reared and infected with *C. albicans* as described, and at 24 hpi larvae were embedded in 0.5% low melting point agarose (LMA) in E3 media plus Tris buffered tricaine methane sulfonate (200 mg/ml) in glass-bottom 24-well imaging dishes. Larvae were scored for recruitment of macrophages and yeast dissemination prior to photoswitching. Photoswitching was done using a 405 nm laser at 10% power on a Olympus IX-81 inverted microscope with an FV-1000 confocal system for 10 minutes on Fluoview XY repeat with a 20X (0.75 NA) objective [13, 29]. After photoswitching, a 10X (0.40 NA) z-stack was taken of the yolk sac area immediately following photoswitching and of the whole fish at 30 and 40 hpi. Imaging was done at room temperature and larvae were kept at 28°C (yeast-locked infections) or 21°C (wild type infections) between imaging sessions. To keep the fish from drying out, 2 mL of E3 media plus PTU was layered over each larva following the first imaging session.

Time lapse imaging for all experiments was done between 32 and 40 hpi unless noted otherwise. For photoswitching experiments, one larva was chosen for time lapse imaging between 32-40 hpi, between 46-54 hpi, and between 56-64 hpi. Larvae were removed from the initial 24-well plate and re-embedded in an 8-well μ-slide insert for the ibidi heating system K-frame (ibidi, Deutschland) for temperature control. The heating system was set up at least one-half hour before starting the time lapse. Time lapses were run on “FreeRun” and images were acquired in succession. Time lapse imaging was done with a 20X (0.75 NA) objective unless noted otherwise. Individual fluorescent channels were compiled on Fluoview software before being transferred for compilation and analysis in Fiji-ImageJ [87].

Fluorescent channels were acquired with optical filters for 635 nm excitation/668 nm emission, 543 nm excitation/572 nm emission, and 488 nm excitation/520 nm emission, for far red fluorescent cells, red fluorescent cells, or green fluorescent cells, respectively. Phalloidin-568 was captured with the optical filter for 543 nm excitation/668 nm emission for histology microscopy. Cascade blue was captured using the optical filter 405 nm excitation/422 nm emission for larvae treated with clodronate liposomes.

### Fiji-ImageJ and MATLAB image analysis

Images were processed in Fiji-ImageJ [87] for counting of phagocytes using Z stack projections (Maximum Intensity Projection), image compilation (Maximum Intensity Projection of the Z stack), and mask compilation. Fiji-ImageJ was also used to track phagocyte movement using the Manual Tracking plugin (Fig. 2). To quantify the amount of TNFα expressing cells or neutrophils in the area of *C. albicans* cells, masks were made of each fluorescent channel (TNFα, neutrophils, and *C. albicans*) in Fiji-ImageJ and run through a script in MATLAB (The MathWorks, Inc., R2017b, Natick, MA). MATLAB workflows are described in Fig. S12. The MATLAB script (Script S1. New_TNF_express) quantified the number of TNFα or neutrophil pixels in the area of *C. albicans* pixels (Figs 4 and 7, Figs. S9 and S11). It also measured the number of fluorescent pixels in the image, so *C. albicans* growth could be measured over time. Note that these calculations were made for individual confocal z-slices and not from maximum projections.

To measure the distance that yeast travelled in Fig. 6D-F, two MATLAB scripts were used. First, distances for each disseminated pixel were measured using (Script S2. Allison_candida_amounts_used). Next, distances were binned by 1-pixel increments and compiled into a single file using the script (Script S3. New_real_Bins_used). Note that these calculations were made for individual confocal z-slices and not from maximum projections. In addition, the data from Allison_candida_amounts_used were also used by the script (Script S4. Allison_combining_candida_amounts_used) to quantify the total level of burden in the yolk and the total amount disseminated. Briefly, masks were made for each fish using the DIC channel to outline the yolk sac and body of the fish. ImageJ masks for the *Candida*, yolk, and body of the fish were read into MATLAB and converted into MATLAB masks for each fish. The amount of *Candida* pixels in the yolk were summed. New *Candida* masks were made to show only disseminated *Candida* pixels and the amount of disseminated *Candida* pixels summed. Distance maps were created from yolk masks giving pixel distances from the yolk. The distance maps and disseminated *Candida* masks were used to find the distances from the yolk for each pixel of *Candida*. The sums of *Candida* in the yolk and disseminated *Candida* as well as the distances for disseminated *Candida* were saved as MATLAB files for each fish to be run through another MATLAB script. The *Candida* sums for the yolk and disseminated *Candida* for each fish were ran through another script to combine the data for each fish into one spreadsheet for *Candida* amounts in the yolk and another for disseminated *Candida* amounts and exported. The *Candida* distances for each fish were run through another script. For each fish the distances were binned so that each bin contained the number of *Candida* pixels at each distance in 1-pixel intervals. The fish were sorted into their respective groups and the bins summed for each group. The groups were then combined into one array and exported into a spreadsheet.

To score and quantify the location (intra- vs. extra-cellular) of disseminated yeast in Rac2-D57N fish treated with clodronate (Fig. 3), 20X images were taken on the Olympus FV1000 confocal at 40 hpi. Fiji-ImageJ was used to score the larvae for yeast that was intra- and extracellular while Photoshop (version CS5 12.1 x64, Adobe Systems Incorporated) was used to trace over yeast scored as intra- and extracellular by methods previously described from our laboratory [5, 13]. PhotoShop layers were annotated and the area calculated in ImageJ.

### Statistical analysis

Statistical analysis was performed in Graphpad Prism versions 6, 9 and 10 (Graphpad software, Inc., La Jolla, CA). All analysis was completed with non-parametric tests (Kruskall-Wallis for ANOVA and Mann-Whitney for pairwise) unless noted otherwise. P-values are indicated throughout the manuscript as: * p≤0.05, ** p≤0.01, *** p≤0.001, **** p≤0.0001. n.s.= not significant.

## Acknowledgements

We would like to thank Brian Peters, Glenn Palmer and Anna Huttenlocher for key *C. albicans* and zebrafish strains and lines. We would like to thank Michel Bagnat for consultation on histology. We would like to thank Julie Walter, Sony Manandhar, Jessica Moore, and Josh Jones for their assistance with initial experiments and data analysis that supported this project. We would like to thank the Henry Lab for advice and sharing. Funding from NIH R15 and R15 (RTW), USDA-Hatch of ME098 (RTW and AKS), Burroughs Wellcome Fund PATH Fellowship (RTW). Funding from NIH fellowship 1F31DK111137-01A1 (JP). RTW is a PATH Investigator of the Burroughs Wellcome Fund.

## Supplemental Figure Legends

**Figure S1. Neither initial *Candida* inoculum nor overall fungal burden indicates whether neutrophils will be recruited to the yolk sac or if the infection will result in dissemination events.** *Tg(mpx:EGFP)* larvae were infected with NRG1^OEX^ as described previously and immediately imaged on the confocal to count the starting inoculum. (A) Example images of larvae just infected (top) and at 40 hpi (bottom), Scale bar = 150 µm. Images demonstrate little difference in starting inoculum resulting in different infection results. (B) CFUs of screened larvae at 0 hpi. Bars show median and interquartile range of yeast per fish. Pooled from three experiments, n = 18 infected fish, median colonies per plate is 19. (C) CFUs from each type of recruitment/dissemination score. Bars indicate median and interquartile range. Pooled from three experiments, left to right n = 20, 8, 6, 20. Stats: Kruskal-Wallis with Dunn’s post-test, no differences with p<0.5.

**Figure S2. Location of *C. albicans* does not affect likelihood of dissemination.** *Tg(mpeg:GAL4)/(UAS:nfsb-mCherry)* fish were crossed with *Tg(fli1:EGFP)* fish and infected with NRG1^OEX^-iRFP as described. Images taken at 24, 30, and 40 hpi were used to quantify the pixel distance of fluorescent *Candida* away from the (A) edge of the sac outlined by the DIC image, or (B) GFP positive endothelial cells lining the vasculature around the yolk sac. Represented are the average distances, for each larva, of each pixel to the closest the yolk edge or EGFP+ cell. Stats: Mann-Whitney tests at each time point. There were no differences with p<0.05. Pooled from 3 experiments, n = 28 non-disseminated larvae and 23 disseminated larvae.

**Figure S3. Killed yeast do not readily disseminate from the yolk sac or elicit immune cell response.** *Tg(mpx:EGFP)* larvae were infected with a wild type *C. albicans* and kept at 21°C for the course of infection. *Candida* was UV killed and stained with AlexaFluor 555 prior to injection in the yolk. Larvae were followed for neutrophil recruitment and fungal dissemination at 24, 30, and 40 hpi. Care was taken to remove larvae with dissemination events that occurred before 24 hpi, to ensure later dissemination events were a result of normal infection processes rather than artifactual injection into the bloodstream. (A) Percent larvae with dissemination of UV inactivated or live fungi at 30 and 40 hpi. For level of dissemination scoring method see Figure 3. Pooled from 4 experiments, left to right n = 77, 85. Stats: Fisher’s exact test, *** p ≤ 0.001, **** p ≤ 0.0001. (B) Percent larvae with neutrophil recruitment to UV inactivated, heat killed, or live fungi. Same larvae as followed in panel A. Stats: Fisher’s exact test, **** p ≤ 0.0001.

**Figure S4. Levels of *tnfa:EGFP* at the site of infection correlate with phagocyte recruitment but not dissemination.** *Tg(lysC:Ds-Red)/Tg(tnfα:GFP)* larvae with red fluorescent neutrophils and green fluorescence with *tnfα* expression were infected with a yeast-locked *C. albicans* as described for Fig. 4. Total GFP+ pixels at 40 hpi were quantified. This was pooled from 5 experiments but was underpowered. N=7, 9 and 5 larvae from L to R. Stats: Kruskal-Wallis with multiple comparisons. p=1.0 and 0.432 respectively for ND/NR vs. ND/R and D/R.

**Figure S5. Overall decrease in the number of phagocytes at the site of infection after dissemination starts.** *Tg(mpeg:GAL4/UAS:nfsb-mCherry)/Tg(mpx:EGFP)* larvae with green fluorescent neutrophils and red fluorescent macrophages were infected as described and scored at 24 hpi. Images at 24, 30 and 40 hpi were quantified as to the number of fluorescent macrophages and neutrophils at the infection site at each time point. Stats: Kruskall-Wallis with Dunn’s post-test. * p<0.05, *** p<0.001. N=12 larvae with original score of Recruited/Disseminated.

**Figure S6. Clodronate liposomes and metronidazole treatment are effective methods for macrophage ablation.** *Tg(mpeg:GAL4/UAS:nfsb-mCherry)/Tg(mpx:EGFP)* larvae with green fluorescent neutrophils and red fluorescent macrophages were used to check efficiency of ablation methods. Larvae were either injected at 28 hpf with 8-10 nl of control or clodronate liposomes (3:1:1 lipsomes:phenol red:10kDa dextran) in the caudal vein or bathed in 20 mM metronidazole for 4 hours following infection and 10 mM metronidazole thereafter. (A) Images of control and clodronate liposome treated larva, scale bar = 150 µm. (B) Number of macrophages or neutrophils counted in a 6 somite region in the trunk at 40 hpi. Bars indicate the median and interquartile range. Liposome treated larvae pooled from 4 experiments, left to right, n = 8, 18, 8, and 18. (C) Images of control and metronidazole treated larvae. (D) Number of macrophages or neutrophils counted in the same region as B. Bars indicate the median and interquartile range. Pooled from 4 experiments, left to right, n = 8, 30, 11, and 19. A one-way ANOVA with Kruskall-Wallis post-test was used to test groups, * ≤ 0.05, ** p ≤ 0.01, *** p ≤ 0.001, **** p ≤ 0.0001, n.s. = not significant.

**Figure S7. Chemical macrophage ablation does not alter dissemination rates.** *Tg(mpeg:GAL4/UAS:nfsb-mCherry)/Tg(mpx:EGFP)* larvae were also used to examine phagocyte recruitment to the infection site. (A) Percent larvae with dissemination in clodronate- (left) and metronidazole- (right) treated larvae. Fisher’s exact test based on numbers shown below in figure, n.s. p>0.05. Pooled from 4 experiments. (B) Infection progression was scored as in Fig. 1, with fish grouped by initial score (top, 24 hpi) and then by final score (bottom, 40 hpi). Fish and experiment numbers are the same as in Panel A.

**Figure S8. Ablation of macrophages by morpholino oligonucleotide does not alter dissemination frequencies.** *Tg(mpeg:GAL4)/(UAS:nfsb-mCherry)/Tg(mpx:EGFP)* embryos were injected at the one-cell stage with a combination of splice blocking and translational blocking *pu.1* morpholino oligonucleotides to inhibit macrophage development. A Cascade blue fluorescent 10 kDa dextran was injected with the morpholino mix, and larvae were screened for correct injection of the morpholino mix following infection with NRG1^OEX^-iRFP. (A-B) Percent larvae with dissemination at 40 hpi. Pooled from 3 independent experiments (n=54 control larvae and n=54 morphant larvae). Stats: Fisher’s exact test, n.s. p>0.05 (C) Representative images of control and ablated larvae at 40 hpi. Images of the anterior and posterior of each fish was stitched with ImageJ, red in yolk is background autofluorescence. Purple outlined boxes are blown up in panel D. Scale bar = 150 µm. (D) Blow-ups of small regions of the head from stitched images in (C) to show loss of macrophages in tissue; the only red fluorescence is from pieces of macrophages that haven’t been cleared. Scale bar = 150 µm.

**Figure S9. Macrophage ablation limits TNFα production without affecting overall burden.** *Tg(LysC:Ds-Red)/Tg(tnfα:GFP)* larvae with red fluorescent neutrophils and green fluorescence with *tnfα* expression were treated with liposomes and infected with NRG1^OEX^-iRFP. (A) Correlation graph between recruited neutrophils and *tnfα* expression overlapping areas of NRG1^OEX^-iRFP fluorescence. (B) ImageJ was used to make masks of the fluorescent channel for the yeast. The number of neutrophil (dsRed) or *tnfα* (GFP) pixels in the area of yeast was measured from these masks in MatLab. Data pooled from 5 experiments, total fish used for quantification, left to right: n= 10, 21, 9, 23. Violin plots. Stats: Mann-Whitney. * p ≤ 0.05. (C) *Candida* burden is not affected by clodronate-mediated macrophage ablation. Total number of *Candida* pixels at the infection site shown as medians and 95% CI. Stats: Mann-Whitney, all p>0.05, not significant.

**Figure S10. Chemical inhibitors of heart beat reduce blood flow, but not phagocyte recruitment or overall yeast dissemination.** *Tg(Mpeg:GAL4/UAS:nfsb-mCherry)/Tg(mpo:EGFP)* larvae were bathed with 2 μM terfenadine/DMSO vehicle or in 0.1 mg/ml valproic acid/E3 water vehicle following infection with NRG1^OEX^-iRFP. (A) Representative images of infected larvae at 40 hpi treated with control or chemical blood flow blockers. (B) Percent area of the infection site with recruited macrophages and neutrophils. Pooled from 3 experiments with terfenadine, left to right, n = 11, 9, 25, and 23. Pooled from 3 experiments with valproic acid, left to right, n = 9, 7, 26, and 15. Stats: two-way ANOVA and Sidak’s multiple comparison’s test, * ≤ 0.05. (C) Dissemination scores of terfenadine treated larvae (pooled from 3 independent experiments) and valproic acid treated larvae (pooled from 3 independent experiments). (D) Details of numbers of individual fish and statistical analysis of experiments shown in Panel C. Fisher’s exact test, n.s. not significant.

**Figure S11. In the context of chemical blockade of blood flow, macrophage ablation reduces *tnfa* expression but doesn’t affect fungal burden.** Rac2-D57N zebrafish were crossed to *Tg(tnfα:GFP)* for neutrophil deficient offspring. All larvae were injected as previously with clodronate liposomes for macrophage ablation, and control or terfenadine for blood flow blockade. Larvae were infected with NRG1^OEX^-iRFP. (A) Representative images of Rac2-D57Nx*Tg(tnfα:GFP)* larvae at 40 hpi. Scale bar = 100 µm. (B) Very low levels of *tnfa* expression in all fish treated with clodronate is unaffected by blockade of blood flow or neutrophil activity. Quantification of *tnfα:GFP* positive pixels in the area of *Candida*, pooled from 5 experiments, left to right n = 13, 25, 12, 23, 13, 24, 12, and 21. Stats: Kruskal-Wallis with Dunn’s post-test, n.s. (C) Total fungal burden as quantified by the number of fluorescent *Candida* pixels in the whole fish (yolk plus body). Pooled from 6 experiments, left to right n = 15, 21, 8, and 10. Stats: Kruskal Wallis with Dunn’s post-test, n.s. Same fish quantified in panels B and C.

**Figure S12. Outline of MATLAB workflows.** This document outlines the steps taken in the MATLAB workflows used for image quantification, referencing the MATLAB scripts that are included as Supplemental Materials.

## MATLAB Script Descriptions

**MATLAB Script S1. New_TNF_express** Quantifies the number of TNFα or neutrophil pixels in the area of *C. albicans* pixels (Figs 4 and 7, Figs. S9 and S11). It also measures the number of fluorescent pixels in the image, so *C. albicans* growth can be measured over time. Note that these calculations were made for individual confocal z-slices and not from maximum projections.

**MATLAB Script S2. Allison_candida_amounts_used**. Measures the shortest distance between each pixel and the yolk edge, for each disseminated pixel, as shown for Fig. 6D-F.

**MATLAB Script S3. New_real_Bins_used.** Bins data from “Allison_candida_amounts_used” by 1-pixel increments and compiles them into a single file. Note that these calculations were made for individual confocal z-slices and not from maximum projections.

**MATLAB Script S4. Allison_combining_candida_amounts_used.** Uses data from the script “Allison_candida_amounts_used” to quantify the total level of burden in the yolk and the total amount of yeast that were disseminated.

## Movie Captions

**Movie S1.** As described in Fig. 2A. *Tg(mpeg1:GAL4/UAS:nfsb-mCherry)/Tg(mpx:EGFP)* larvae were infected with yeast-locked *C. albicans* and scored for recruitment of phagocytes and dissemination of yeast at 24 hpi. A time-lapse of a larva with recruitment was taken over the course of 7 hours starting at ∼32 hpi. Neutrophils and macrophages were observed moving within and away from the infection site carrying yeast.

**Movie S2.** As described in Fig. 2A, but from an independent experiment. *Tg(mpeg1:GAL4/UAS:nfsb-mCherry)/Tg(mpx:EGFP)* larvae were infected with yeast-locked *C. albicans* and scored for recruitment of phagocytes and dissemination of yeast at 24 hpi. A time-lapse of fish with recruitment was taken over the course of 7 hours starting at ∼32 hpi. Neutrophils and macrophages were observed moving within and away from the infection site carrying yeast.

**Movie S3.** As described in Fig. 2C. *Tg(mpeg1:GAL4)x(UAS:Kaede)* larvae were infected with yeast-locked *C. albicans* and scored for recruitment of macrophages and dissemination of yeast at 24 hpi. Macrophages near the infection site of larvae with immune recruitment were photo-switched at 24 hpi. A time-lapse movie was taken over the course of 7 hours starting at ∼46 hpi. A photo-converted macrophage moves out of the blood stream into tail tissue.

**Movie S4.** As described in Fig. 2D. *Tg(mpeg1:GAL4)x(UAS:Kaede)* larvae were infected with yeast-locked *C. albicans* and scored for recruitment of macrophages and dissemination of yeast at 24 hpi. Macrophages near the infection site of larvae with immune recruitment were photo-switched at 24 hpi. A time-lapse movie taken over the course of 7 hours starting at ∼32 hpi. Intracellular yeast growth and apparent NLE events are highlighted.

**Movie S5.** *Tg(mpx:EGFP)* larvae were infected with hypofilamentous *C. albicans* strain *cph1*Δ/Δ*efg1*Δ/Δ-dTomato in the yolk as described and imaged at 44 hpi by time-lapse microscopy. Neutrophils (green) interacting with yeast (red) as they move down the tail tissue inside a blood vessel.

**Movie S6.** As described in Fig. 4D. *Tg(tnfα:EGFP)* larvae with green fluorescence with *tnfα* expression were given control liposomes and infected with a yeast-locked *C. albicans* as described. Z series demonstrating typical *tnfα* expression at the infection site in a wild type larva.

**Movie S7.** As described in Fig. 4D. *Tg(tnfα:EGFP)* larvae with green fluorescence with *tnfα* expression were given clodronate liposomes and infected with a yeast-locked *C. albicans* as described. Z series demonstrating background *tnfα* expression in a wild type larva.

**Movie S8.** As described in Fig. 4E. Rac2-D57N/*Tg(tnfα:EGFP)* larvae with red fluorescent neutrophils and green fluorescence with *tnfα* expression were given control liposomes and infected with a yeast-locked *C. albicans* as described. Z series demonstrating typical *tnfα* expression at the infection site in a Rac2/D57N larva.

**Movie S9.** As described in Fig. 4E. Rac2-D57N/*Tg(tnfα:EGFP)* larvae with red fluorescent neutrophils and green fluorescence with *tnfα* expression were given clodronate liposomes and infected with a yeast-locked *C. albicans* as described. Z series demonstrating background *tnfα* expression in a Rac2/D57N larva.

**Movie S10.** As described in Fig. 4. Rac2-D57N/*Tg(tnfα:EGFP)* larvae with red fluorescent neutrophils and green fluorescence with *tnfα* expression were given control liposomes and infected with a yeast-locked *C. albicans* as described. Time-lapse movie taken over the course of 7 hours starting at ∼32 hpi. Macrophage-like cells turn on *tnfa* expression as they move closer to the site of infection.

**Movie S11.** As described in Fig. 4. Rac2-D57N/*Tg(tnfα*:*EGFP)* larvae with red fluorescent neutrophils and green fluorescence with *tnfα* expression were given clodronate liposomes and infected with a yeast-locked *C. albicans* as described. Time-lapse movie taken over the course of 7 hours starting at ∼32 hpi. There is an absence of phagocytes turning on *tnfa* expression.

**Movie S12.** As described in Fig. S10. Tg*(mpeg:GAL4/UAS:nfsb-mCherry)/Tg(mp*x*:EGFP)* larvae were bathed with 2 μM terfenadine/DMSO vehicle following infection with NRG1^OEX^-iRFP. Representative movie of infected larvae at ∼32 hpi treated with 2 μM terfenadine, which blocks blood flow.

**Movie S13**. As described in Fig. S10. *Tg(mpeg:GAL4/UAS:nfsb-mCherry)/Tg*(*mp*x*:EGFP)* larvae were bathed in 0.1 mg/ml valproic acid/E3 water vehicle following infection with NRG1^OEX^-iRFP. Representative movie of infected larvae at ∼32 hpi treated with 0.1 mg/ml valproic acid, which blocks blood flow.

**Movie S14.** As described in Fig. 5G. *Tg(fli1:EGFP)* larvae, with GFP-expressing endothelial cells, were infected with *NRG1*^OEX^-iRFP as previously described. A vehicle-treated *Tg(fli1:EGFP)* fish with intact blood flow was imaged at ∼32 hpi by time-lapse on the confocal.

**Movie S15.** As described in Fig. 5G. *Tg(fli1:EGFP)* larvae, with GFP-expressing endothelial cells, were infected with *NRG1*^OEX^-iRFP as previously described. A vehicle-treated *Tg(fli1:EGFP)* fish with intact blood flow was imaged at ∼32 hpi by time-lapse on the confocal.

**Movie S16.** As described in Fig. 7. RAC2-D57N/*Tg(tnfα:GFP)* larvae were injected with control liposomes and infected with the wild type CAF2-iRFP *C. albicans* as described for the yeast-locked infections. Fish were imaged by confocal microscopy between 32 and 40 hpi.

**Movie S17.** As described in Fig. 7. RAC2-D57N/*Tg(tnfα:GFP)* larvae were injected with clodronate liposomes and infected with the wild type CAF2-iRFP *C. albicans* as described for the yeast-locked infections. Fish were imaged by confocal microscopy between 32 and 40 hpi.

**Movie S18.** As described in Fig. 7. *Tg(mpeg:GAL4/UAS:nfsb-mCherry)/Tg*(*mp*x*:EGFP)* larvae were infected with the wild type Caf2-iRFP *C. albicans* as described for the yeast-locked infections. Fish were imaged by confocal microscopy between 32 and 40 hpi.

